# Fitness and immune-escape within germinal centers shape premalignant evolution toward lymphoma

**DOI:** 10.64898/2026.06.25.734549

**Authors:** Lingling Zhang, Miu Shing Hung, Oscar Atkins, Pavel Artemov, Akhesa Sochon, Victoire Boulat, Hamid Kashkar, Hans Christian Reinhardt, Jude Fitzgibbon, Jessica Okosun, Dinis Pedro Calado

**Affiliations:** Immunity and Cancer Laboratory, The Francis Crick Institute, UK; Immunity and Cancer Laboratory, The Babraham Institute, Babraham Research Campus, Cambridge, UK; Immunology Programme, The Babraham Institute, Babraham Research Campus, Cambridge, UK; Centre for Haemato-Oncology, Barts Cancer Institute, Queen Mary University of London, London, UK; University Hospital Cologne, Germany; Department of Hematology and Stem Cell Transplantation, University Hospital Essen, German; Centre for Cancer Evolution, Barts Cancer Institute, Queen Mary University of London, London, UK

**Keywords:** Germinal center immunosurveillance, premalignant GC B-cells, follicular lymphoma, in situ follicular neoplasia, CD8⁺ T-cell surveillance, immune escape, B-cell fitness, lymphoma evolution, cancer immunoediting, germinal center microenvironment

## Abstract

Germinal centers (GCs) support physiological B-cell mutagenesis and are considered lymphoma-permissive; nevertheless, lymphoma development is uncommon. Human *in situ* follicular neoplasia (ISFN) captures this paradox: premalignant B-cells can persist within GCs for prolonged periods without progressing to overt lymphoma. We found that human ISFN, but not normal GCs, are infiltrated by CD8⁺ T-cells, suggesting that premalignant GC B-cells are locally immune-surveilled. Using mouse models that separate early premalignant fitness from lymphoma-associated evolution, we show that fitness-enhanced premalignant GC B-cells expand transiently but are selectively eliminated by infiltrating cytotoxic CD8⁺ T-cells, while normal GC B-cells are spared. By contrast, evolved premalignant GC B-cells retain their fitness but disable productive CD8⁺ T-cell cytotoxic differentiation, allowing persistence and lymphoma-like transcriptional and genomic evolution. These findings establish GCs as active immune-surveillance sites and show that progression from premalignancy to lymphoma requires both enhanced GC fitness and escape from local immune control.

**Key findings:**

GCs undergo active immune-surveillance to detect premalignant B-cells.
Premalignant GC B-cells trigger cytotoxic CD8⁺ T-cell responses.
Lymphoma-associated evolution enables immune-escape within GCs.
Fitness and immune-escape drive evolution from premalignancy to lymphoma.

**Blurb:** Germinal centers are considered lymphoma-permissive; however, progression from premalignancy is uncommon. Using models of human *in situ* follicular neoplasia, Zhang et al. demonstrate that infiltrating CD8⁺ T-cells actively eliminate premalignant GC B-cells. Co-occurrence of lymphoma-like alterations blocks this cytotoxic T-cell response, driving immune escape and lymphoma evolution.

## Introduction

Cancer evolution is usually reconstructed from established tumors; nevertheless, the selective pressures that determine whether premalignant cells progress operates much earlier, within organized tissues^1–5^. Germinal centers (GCs) provide an unusually tractable setting to study this process. GC B-cells physiologically mutate their DNA to generate high-affinity antibodies^6–11^, creating repeated opportunities for oncogenic alterations. Consequently, GCs B-cells give rise to most B-cell lymphomas^12–14^. The paradox is that premalignant GC B-cells are common, whereas progression to lymphoma is uncommon^15–18^. Founder alterations alone appear insufficient for progression to lymphoma, implying that premalignant GC B-cells must survive local selective pressures within GC reactions.

We asked whether premalignant B-cell evolution toward lymphoma requires not only acquisition of GC fitness, enabling mutant B-cells to expand, but also to immune-escape from local surveillance triggered by their altered state. Consistent with this possibility, defects in perforin-mediated cytotoxicity predispose to lymphoma^19–24^. However, whether such immune-surveillance operates at the premalignant stage within organized GCs remains unresolved. GCs are considered immune-privileged, lymphoma-supportive environments shaped by Tfh-and stromal-cell interactions^25–33^. This raises a central question: can cytotoxic cells enter premalignant GCs and eliminate altered B-cells before lymphoma develops?

Human *in situ* follicular neoplasia (ISFN) provides a direct window into this paradox^34^. ISFN is defined by GC-constrained clonal BCL2⁺ B-cells, often carrying *CREBBP* alterations^35–37^. Although detected incidentally in up to 4% of individuals, ISFN progression to lymphoma is uncommon^15–18^. We found that ISFN GCs, unlike normal GCs, contained CD8⁺ T-cells, suggesting that premalignant GC B-cells are subject to local immune-surveillance.

To test this experimentally, we engineered mice with contrasting alterations associated with ISFN versus lymphoma. *Bcl2-*overexpression, with or without *Crebbp*-loss, modeled early premalignant GC B-cell states associated with ISFN. Because *KMT2D*-mutations are enriched in overt lymphoma and are the most frequent co-occurrence with *BCL2/*CREBBP founding alterations^38–40^, we added *Kmt2d-*loss to model an evolved state. This genetic staging revealed that *Bcl2*/*Crebbp*-altered GC B-cells acquired increased fitness but were immune-surveilled: cytotoxic CD8⁺ T-cells infiltrated GCs and eliminated these mutant cells while sparing normal GC B-cells in the same mouse. By contrast, *Kmt2d*-loss preserved GC B-cell fitness but uncoupled CD8⁺ T-cell infiltration from productive killing by blocking terminal cytotoxic differentiation, a defect reversed by TGF-β receptor blockade. This immune-escape enabled lymphoma-like transcriptional and genomic evolution.

Thus, GCs are not immune-privileged sites but actively surveilled environments, where lymphoma evolution depends on both premalignant GC B-cell fitness and escape from CD8⁺ T-cell control.

## Results

### Premalignant GC B-cells are fitness-enhanced but immune-surveilled

If premalignant GC B-cells are constrained by local immune-surveillance, human ISFN should show evidence of immune-cell recruitment with cytotoxic capacity. To test this, we performed multiplex immunofluorescence on lymph node biopsies from individuals with ISFN and on control lymph nodes. CD20⁺ CD10⁺ BCL6⁺ GCs were readily identified in both settings, whereas ISFN GCs showed enrichment of BCL2⁺ GC B-cells^41,42^, consistent with their premalignant biology (**Fig.1A**,**B**). GCs from control lymph nodes contained few CD8⁺ T-cells. By contrast, ISFN GCs showed prominent CD8⁺ T-cell infiltration (**Fig.1A**,**B**). NCAM1⁺ NK-cells were rare or absent in both settings, indicating that CD8⁺ T-cells dominated the infiltrate within ISFN GCs (**Fig.1A**,**B**). These findings suggest that ISFN GCs are locally immune-surveilled, with CD8⁺ T-cells representing the dominant population recruited to premalignant ISFN GCs.

**Figure 1.**
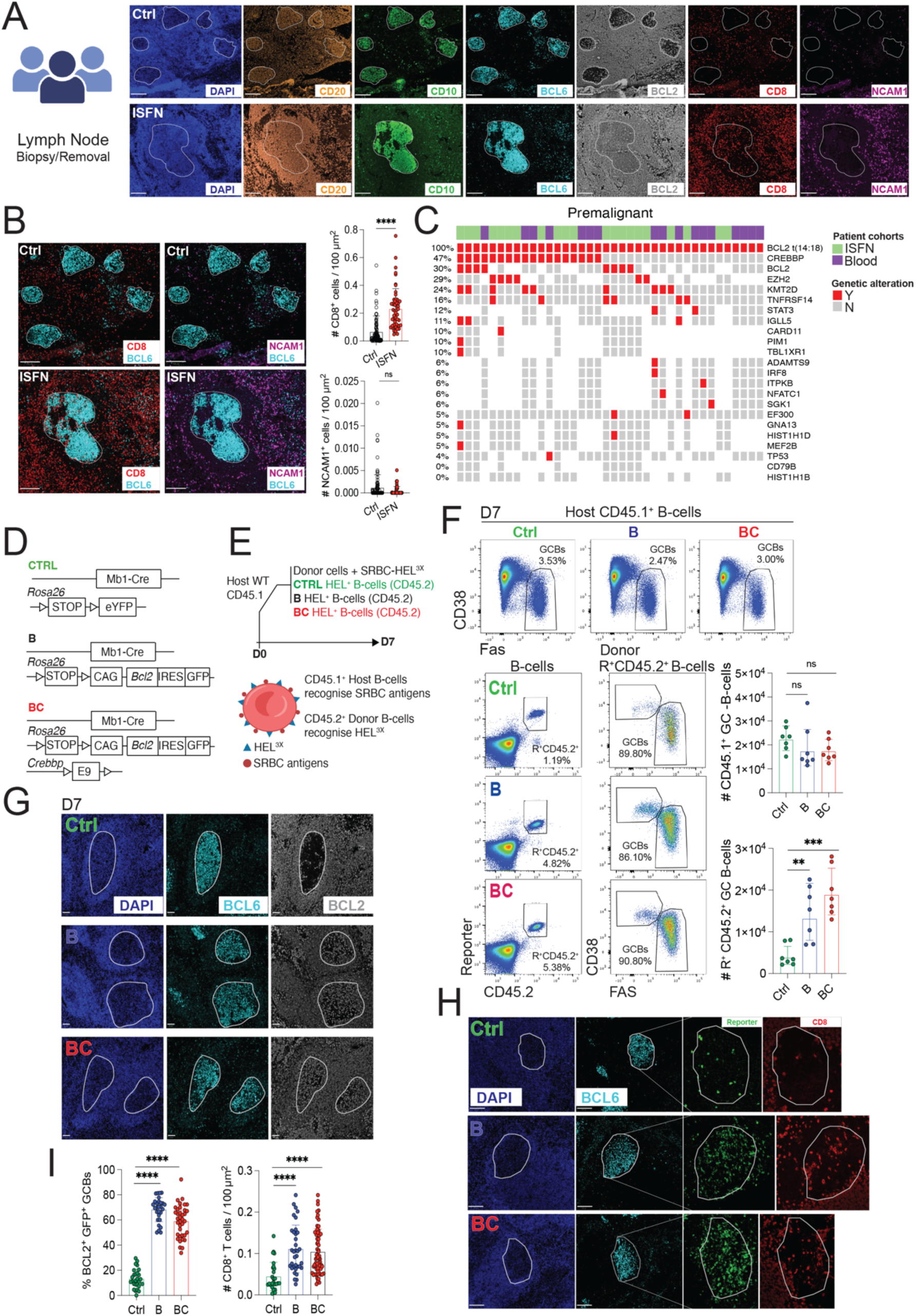
Premalignant GC B-cells are fitness-enhanced but immune-surveilled. **(A-B)** Multiplex immunofluorescence analysis of control lymph nodes and lymph nodes from individuals with *in situ* follicular neoplasia (ISFN). **(A)** Representative staining showing DAPI (blue), CD20 (orange), CD10 (green), BCL6 (cyan), BCL2 (grey), CD8 (red), and NCAM1 (purple). Scale bar, 200 μm. **(B)** Left: representative counterstains highlighting BCL6⁺ GCs with CD8⁺ T-cells (left; BCL6, cyan; CD8, red) or NCAM1⁺ NK-cells (right; BCL6, cyan; NCAM1, purple). Scale bar, 200 μm. Right: quantification of CD8⁺ T-cells and NCAM1⁺ NK-cells within GCs from control and ISFN lymph nodes. **(C)** Recurrent genetic alterations in lymphoma premalignancy, integrating exome-sequencing data from ISFN samples (green) and pre-diagnostic peripheral blood samples (purple). **(D)** Schematic of allele combination in control (Ctrl), *Bcl2* overexpression (B) and *Bcl2* overexpression with heterozygous *Crebbp* loss (BC) mice in the SWHEL system. **(E)** Top: experimental design for adoptive transfer experiment. Bottom: schematic showing the two concurrent GC responses induced in recipient mice by SRBC-HEL^3x^ immunization: an endogenous CD45.1⁺ response to SRBCs and a donor-derived CD45.2⁺ SWHEL response to HEL^3x^. **(F)** GC B-cell responses 7 days after immunization. Top: Endogenous CD45.1⁺ GC B-cell response to SRBCs. Bottom: Left, representative gating strategy for reporter^+^ CD45.2⁺ donor derived B-cells and GC B-cells; right, quantification of endogenous CD45.1⁺ GC B-cell numbers per 10⁶ splenocytes and for donor-derived reporter^+^ CD45.2⁺ GC B-cells per 10⁶ splenocytes. **(G-I)** Immunofluorescence analysis of spleen sections from immunized recipient mice 7 days after transfer of Ctrl, B, or BC donor B-cells. **(G)** Representative staining showing DAPI (blue), BCL6 (cyan), BCL2 (grey). Scale bar, 50 μm. **(H)** Detection of CD8⁺ T-cells within GCs. Left, representative counterstaining for BCL6 (cyan), reporter (green) and CD8 (red). Scale bar, 100 μm. **(I)** Left, quantification of the percentage of BCL2⁺ cells among reporter⁺ GC B-cells. Bottom, quantification of CD8⁺ T-cells per GC. Each symbol in **(B, I)** represents an individual GC. Each symbol in **(F)** represents an individual mouse. **(A)** control n = 7 and ISFN n = 6 samples; **(F)** Ctrl n = 7, B n = 7, and BC n = 7 mice; **(G, H and I)** Ctrl n = 3, B n = 3, and BC n = 3 mice. Small horizontal lines indicate mean ± SD. Data in **(F)** are from two independent experiments, Data in **(G, H and I)** are data representative from three biological replicates. **P ≤ 0.01, ***P ≤ 0.001, ****P ≤ 0.0001; unpaired two-tailed Student’s t test. ns, not significant.

To determine whether this observation in the human could be modeled experimentally, we engineered mice contrasting alterations associated with either ISFN-like premalignancy or lymphoma evolution. Genetic analyses of ISFN and premalignant B-cell states in healthy individuals identify *BCL2*t(14:18) as a founding event, present in 100% of cases, with additional *CREBBP*-mutations observed in 47% of cases^35–37^ (**Fig.1C**). We therefore used *Bcl2*-overexpression, alone or together with *Crebbp*-loss, to generate tractable ISFN-like premalignant GC B-cell states.

We crossed *Cd79a-cre* mice^43^ with a conditional *R26*-*Bcl2*^LSL^ allele^44^ to enforce *Bcl2* expression, generating *BCL2*-overexpressing mice, hereafter referred to as B mice (**Fig.1D**). The timing of *Cd79a-cre* activity coincides with RAG-mediated recombination, providing a developmental context analogous to the emergence of *BCL2*t(14;18) translocations^45^. We then combined *R26*-*BCL2*^LSL^ with heterozygous *Crebbp*-loss^35,46,47^ to generate *BCL2-*overexpressing/*Crebbp*-deficient mice, hereafter referred to as BC mice (**Fig.1D**). In both B and BC mice, Cre-recombined B-cells were marked by GFP expression from the *R26-BCL2^LSL^* allele. Control mice, hereafter referred to as Ctrl mice, carried a *Cd79a-cre* together with the *R26-YFP^LSL^* reporter allele^48^. Incorporation of the SWHEL B-cell receptor knock-in system^49^ enabled controlled recruitment of HEL-specific CD45.2⁺ donor B-cells into GC responses after transfer into wild-type CD45.1⁺ recipients and immunization with SRBC-HEL^3X^ (**Fig.1E**). Transfer of small numbers of B, and BC donor B-cells recapitulated the sporadic nature of premalignant B-cells^12^ and generated two concurrent GC responses in the same mouse: an endogenous CD45.1⁺ response to SRBCs and a donor-derived CD45.2⁺ reporter⁺ response to HEL3x (**Fig.1E**,**S1A**).

Host-derived CD45.1⁺ endogenous GC responses were comparable across recipients, irrespective of the genotype of the transferred donor B-cells, indicating that the recipient environment supported physiological GC formation (**Fig.1F**). In contrast, donor-derived *BCL2*-overexpressing B-cells (B) and *BCL2*-overexpressing/*Crebbp-*deficient B-cells (BC) expanded within GCs and displayed features of premalignant GC hyperplasia, consistent with previous mouse models^42^ and human ISFN GCs (**Fig.1A**). Supporting an ISFN-like phenotype, BCL2⁺ B-cells accumulated significantly within GCs of immunized recipient mice receiving B or BC B-cells, but not Ctrl B-cells (**Fig.1G**). Together, these data show that ISFN-like GC states in mice are driven by enhanced premalignant GC B-cell fitness, reflected by expansion of donor-derived B and BC cells and their increased representation within GC structures, compared to Ctrl B-cells (**Fig.1F-I**).

Having recapitulated key features of human ISFN GCs (**Fig.1A**,**B**), we next asked whether mouse premalignant GCs similarly recruit immune cells. We therefore stained for CD8⁺ T-cells and NCAM1⁺ NK-cells (**Fig.1H**,**S1B**). Premalignant GCs formed by donor-derived B or BC B-cells showed prominent CD8⁺ T-cell infiltration compared with GCs formed by Ctrl B-cells (**Fig.1H**, **I**). By contrast, NK-cells were not detected in either setting (**Fig.S1B**). Thus, as in human ISFN, mouse premalignant GCs selectively recruited CD8⁺ T cells rather than NK cells.

Together, these data show that premalignant B-cells carrying ISFN-associated alterations are fitter than normal GC B-cells and can expand within GC reactions. However, this enhanced fitness is accompanied by local immune-surveillance, marked by CD8⁺ T-cell infiltration into premalignant GCs. This human-to-mouse convergence established a tractable system to test whether CD8⁺ T-cell infiltration is a bystander feature of premalignancy or an active barrier to lymphoma evolution.

### CD8⁺ T-cells selectively eliminate premalignant GC B-cells

Having shown that ISFN-like premalignant GC B-cells acquire increased GC fitness but are subject to immune-surveillance, we next asked whether CD8⁺ T-cells actively eliminate them.

We first followed the dynamics of donor-derived B-cells after GC formation. Compared with Ctrl donor B-cells, B and BC B-cells showed an early expansion advantage at day 7 after immunization, consistent with increased premalignant GC fitness (**Fig.1F**). However, this advantage was transient. As the response progressed, B- and BC-derived GC B-cells declined sharply at days 9 and 11 after immunization (**Fig.2A-C**). By day 11, B and BC donor-derived premalignant GC B-cells were nearly lost, whereas Ctrl donor-derived GC B-cells remained detectable and host endogenous CD45.1⁺ GC B-cells were preserved (**Fig.2B-D**,**S2A**).

**Figure 2.**
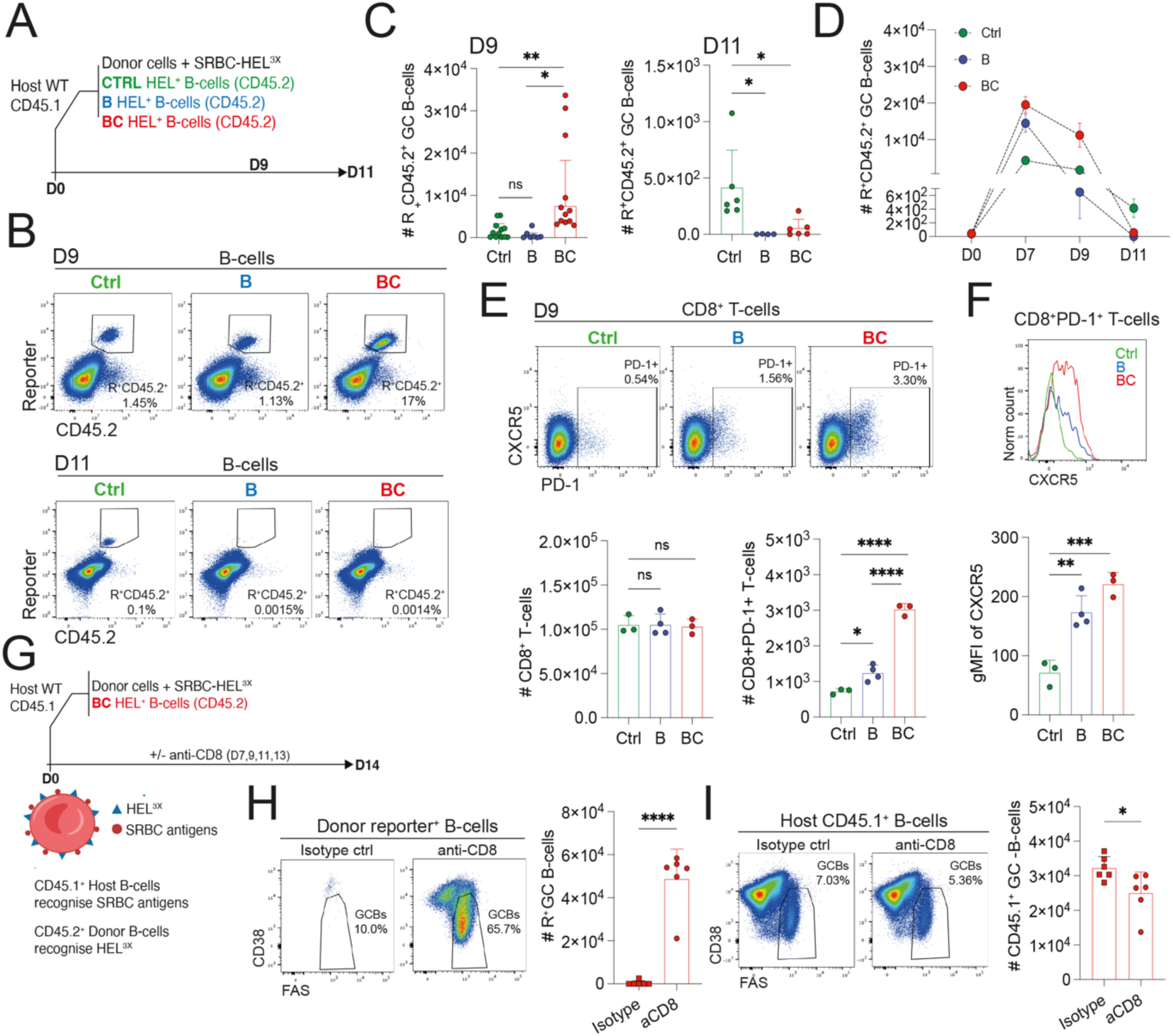
CD8⁺ T-cells selectively eliminate premalignant GC B-cells. **(A)** Experimental design for tracking donor-derived Ctrl, B, and BC B-cell responses after transfer into CD45.1⁺ recipient mice and immunization with SRBC-HEL^3x^. **(B-C)** Donor-derived CD45.2⁺ B-cell responses 9 and 11 days after immunization. **(B)** Top, representative gating of donor-derived GFP⁺CD45.2⁺ B-cells on day 9. Bottom, representative gating of donor-derived CD45.2⁺ GC B-cells on day 11. **(C)** Quantification of donor-derived GC B-cell numbers per 10⁶ splenocytes. **(D)** Summary of donor-derived Ctrl, B, and BC GC B-cell dynamics during the GC response. GC B-cell numbers per 10⁶ splenocytes. **(E)** CD8⁺ T-cell activation in recipient mice. Top, representative gating of PD-1⁺ cells among CD8⁺ T-cells. Bottom, quantification of total CD8⁺ T-cells and PD-1⁺ CD8⁺ T-cells per 10⁶ splenocytes. **(F)** CXCR5 expression by PD-1⁺ CD8⁺ T-cells. Top, representative histograms of CXCR5 expression across conditions. Bottom, quantification of CXCR5 geometric mean fluorescence intensity (gMFI) on PD-1⁺CD8⁺ T-cells. **(G)** Experimental design for CD8⁺ T-cell depletion after GC establishment. **(H)** Donor-derived BC GC B-cell responses 14 days after immunization in mice treated with isotype control or anti-CD8 antibody. Representative gating and quantification of reporter⁺CD45.2⁺ donor GC B-cells per 10⁶ splenocytes. **(I)** Endogenous CD45.1⁺ GC B-cell responses 14 days after immunization in mice treated with isotype control or anti-CD8 antibody. Representative gating of CD45.1⁺ GC B-cells and quantification of endogenous GC B-cell numbers per 10⁶ splenocytes. Each symbol in **(C, E, F, H, I)** represents an individual mouse. **(C)** D9: Ctrl n = 12, B n = 7, BC n = 12. D11: Ctrl n = 6, B n = 4, BC n = 6. **(E-F)** Ctrl n = 3, B n = 4, BC n = 3. **(H-I)** isotype n = 6, anti-CD8 n = 6. Small horizontal lines indicate mean ± SEM. Data in **(C)** are from three independent experiments; data in **(E-F)** are representative of three independent experiments; data in **(H-I)** are representative of two independent experiments; *P ≤ 0.05, **P ≤ 0.01, ***P ≤ 0.001, ****P ≤ 0.0001; unpaired two-tailed Student’s t test. ns, not significant.

We next tested whether CD8^+^ T-cells accounted for the disappearance of premalignant GC B-cells. CD8⁺ T-cells elicited by B- and BC-derived premalignant GC B-cells displayed features of local GC immune-surveillance. Total CD8⁺ T-cell numbers were similar across conditions, but recipients of B and BC B-cells contained increased numbers of PD-1⁺ CD8⁺ T-cells (**Fig.2E**). These activated PD-1⁺ CD8⁺ T-cells expressed elevated CXCR5, consistent with the acquisition of GC-homing capacity^50,51^ (**Fig.2F**).

BC-derived premalignant GC B-cells showed the most robust early expansion followed by the most pronounced disappearance (**Fig.1F**,**2B-D**). We therefore transferred BC B-cells, allowed premalignant GCs to establish, and then depleted CD8^+^ T-cells using an anti-CD8 antibody from day 7 onwards (**Fig.2G**). CD8^+^ T-cells depletion was highly efficient (**Fig.S2B**) and rescued BC-derived premalignant GC B-cell expansion (**Fig.2H**). This rescue enhanced the competitive representation of BC-derived premalignant GC B-cells relative to host-derived endogenous CD45.1^+^ GC B-cells (**Fig.2I**). Thus, the disappearance of fitness-enhanced premalignant GC B-cells is not passive loss but active CD8⁺ T-cell-mediated elimination.

Together, these findings show that premalignant GC B-cells trigger the recruitment and activation of CD8⁺ T-cells, resulting in selective elimination of fitness-enhanced mutant B-cells. Thus, GC fitness is conditional: premalignant GC B-cells can expand, but persistence requires survival of CD8⁺ T-cell surveillance. This may explain why human ISFN is detectable as a premalignant GC state, but only rarely progresses to overt lymphoma.

### Lymphoma-associated evolution enables immune-escape within GCs

Because ISFN-like premalignant GC B-cells were eliminated by CD8⁺ T-cells, we asked whether lymphoma-enriched alterations convert immune-surveilled mutant B-cells into immune-evasive. We focused on *KMT2D* loss because mutations in *KMT2D* are much more highly recurrent in overt follicular lymphoma (>70% of tumors) compared to premalignant samples (24%) and represent the most frequent co-occurring alteration on a *BCL2*/*CREBBP*-altered genetic background^38–40^ (**Fig.1C** and **Fig. 3A**).

**Figure 3.**
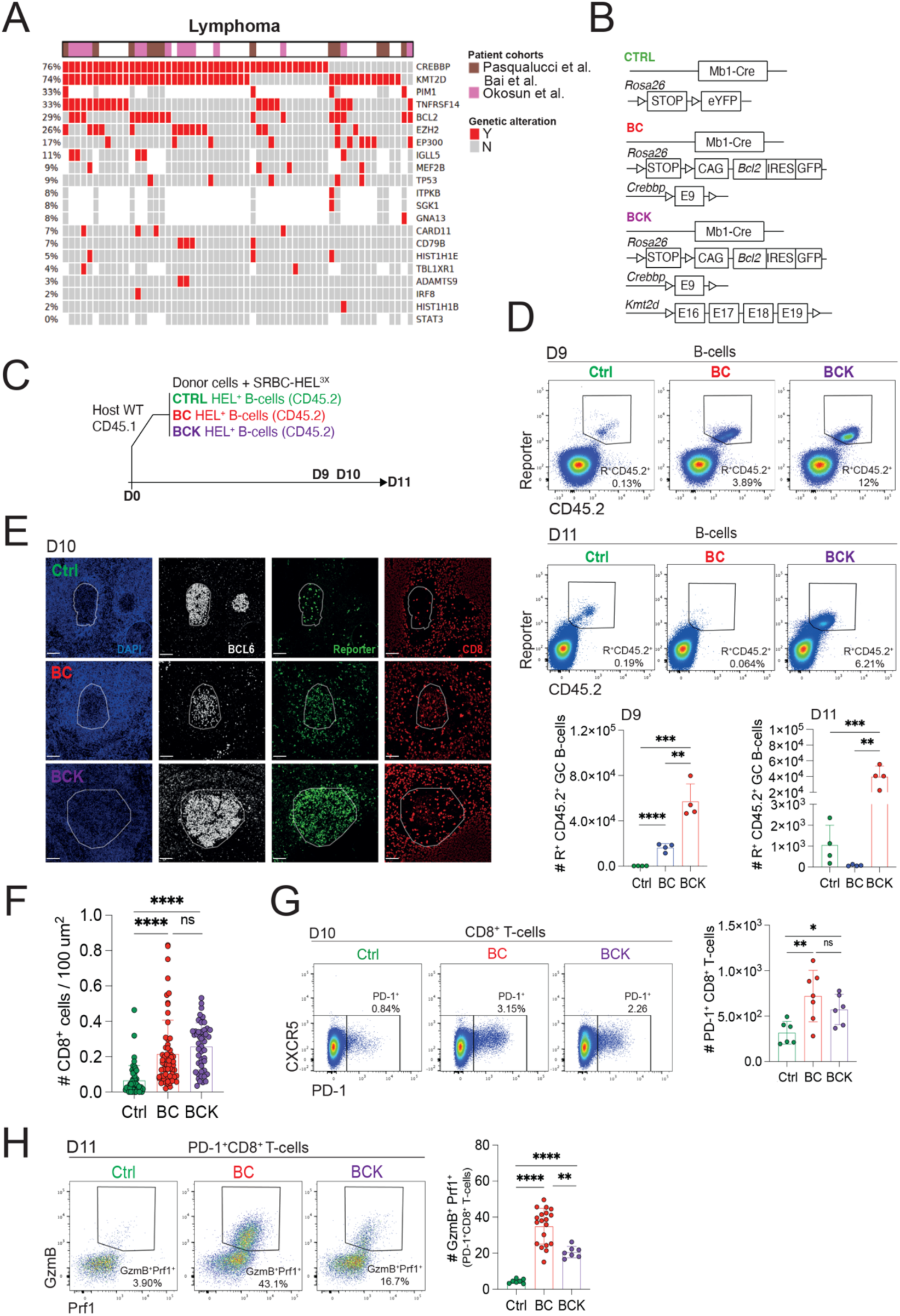
Lymphoma-associated evolution enables immune-escape within GCs. **(A)** Recurrent genetic alterations in overt lymphoma, based on exome-sequencing data from follicular lymphoma patient samples. **(B)** Schematic of allele combination in control (Ctrl), *Bcl2* overexpression with heterozygous *Crebbp* loss (BC) and *Bcl2*-overexpressing mice with heterozygous *Crebbp* loss combined with *Kmt2d* loss (BCK) mice in the SWHEL system. **(C)** Top: experimental design for adoptive transfer experiment. **(D)** Donor-derived CD45.2⁺ B-cell responses 9 and 11 days after immunization. Top, representative gating of donor-derived reporter⁺ CD45.2⁺ B-cells on day 9. Middle, representative gating of donor-derived reporter⁺ CD45.2⁺B-cells on day 11. Bottom, quantification of donor-derived GC B-cell numbers per 10⁶ splenocytes. **(E)** CD8⁺ T-cell infiltration into GCs. Left, representative immunofluorescence staining of spleen sections showing DAPI (blue), BCL6 (cyan), reporter⁺ donor cells (green), and CD8 (red). Scale bar, 50 μm. **(F)** Quantification of data in (E) for CD8⁺ T-cells per GC. **(G)** Activated CD8⁺ T-cell responses in recipient mice. Left, representative gating of PD-1⁺ cells among CD8⁺ T-cells. Right, quantification of PD-1⁺CD8⁺ T-cells per 10⁶ splenocytes. **(H)** Left, representative gating of GranzymeB⁺ perforin⁺ effector cells among PD-1⁺CD8⁺ T-cells at day 11 after immunization. Right, quantification of GranzymeB⁺ perforin⁺ effector cells among PD-1⁺ CD8⁺ T-cells. Each symbol in **(D, G, H)** represents an individual mouse; Each symbol in **(F)** represents an individual GC. **(D)** D9: Ctrl n = 4, BC n = 4, BCK n = 4; D11: Ctrl n = 4, BC n = 4, BCK n = 4. **(E)** Ctrl n = 3, BC n = 3, BCK n = 3; **(G)** Ctrl n = 6, BC n = 7, BCK n = 6. **(H)** day 11: Ctrl n = 7, BC n = 19, BCK n = 7. Small horizontal lines indicate mean ± SD. Data in **(D)** are representative of three independent experiments; data in **(G)** are from two independent experiments; data in **(H)** are representative of three independent experiments; and *P ≤ 0.05, **P ≤ 0.01, ***P ≤ 0.001, ****P ≤ 0.0001; unpaired two-tailed Student’s t test. ns, not significant.

We therefore hypothesized that *KMT2D*-loss defines a progression-associated state in which premalignant GC B-cells become resistant to immune-surveillance. To model this state, we introduced *Kmt2d-*deletion^52^ into the immune-surveilled BC model, generating BCK mice (**Fig.3B**). Donor BCK B-cells were transferred and tracked after immunization (**Fig.3C**). BCK-derived premalignant GC B-cells behaved distinctively from immune-surveilled BC B-cells. At day 9 after immunization, BCK-derived GC B-cells showed greater expansion than BC-derived B-cells (**Fig.3D**). By day 11, when BC-derived premalignant GC B-cells had collapsed, BCK-did not (**Fig.3D**).

We asked how BCK-derived premalignant GC B-cells escaped CD8⁺ T-cell immune-surveillance. One possibility was that *Kmt2d*-loss prevented CD8⁺ T-cell recruitment or activation; however, this was not the case. Both BC and BCK-derived premalignant GCs showed elevated CD8⁺ T-cell infiltration and increased numbers of PD-1⁺ CD8⁺ T-cells compared with Ctrl-derived GCs (**Fig.3E-G**). Thus, *Kmt2d-*loss did not abolish immune-recognition but altered the functional outcome of CD8⁺ T-cell activation.

Indeed, BC B-cell recipients accumulated granzyme-B⁺ perforin-1⁺ PD-1⁺ CD8⁺ T-cells, coinciding with elimination of BC-derived premalignant GC B-cells (**Fig. 3H**). In contrast, BCK B-cell recipients showed CD8⁺ T-cell activation and entry into premalignant GCs but failed to sustain a cytotoxic effector phenotype (**Fig.3H**).

Thus, lymphoma-associated evolution preserves premalignant GC B-cell fitness while disabling the cytotoxic arm of CD8⁺ T-cell surveillance.

### Immune-escape arises from impaired CD8⁺ T-cell effector differentiation

To define how evolved premalignant GC B-cells alter CD8⁺ T-cell fate, we sorted PD-1⁺ CD8⁺ T-cells from recipient mice 10 days after immunization and performed single-cell RNA sequencing (scRNA-seq) (**Fig.4A**).

**Figure 4.**
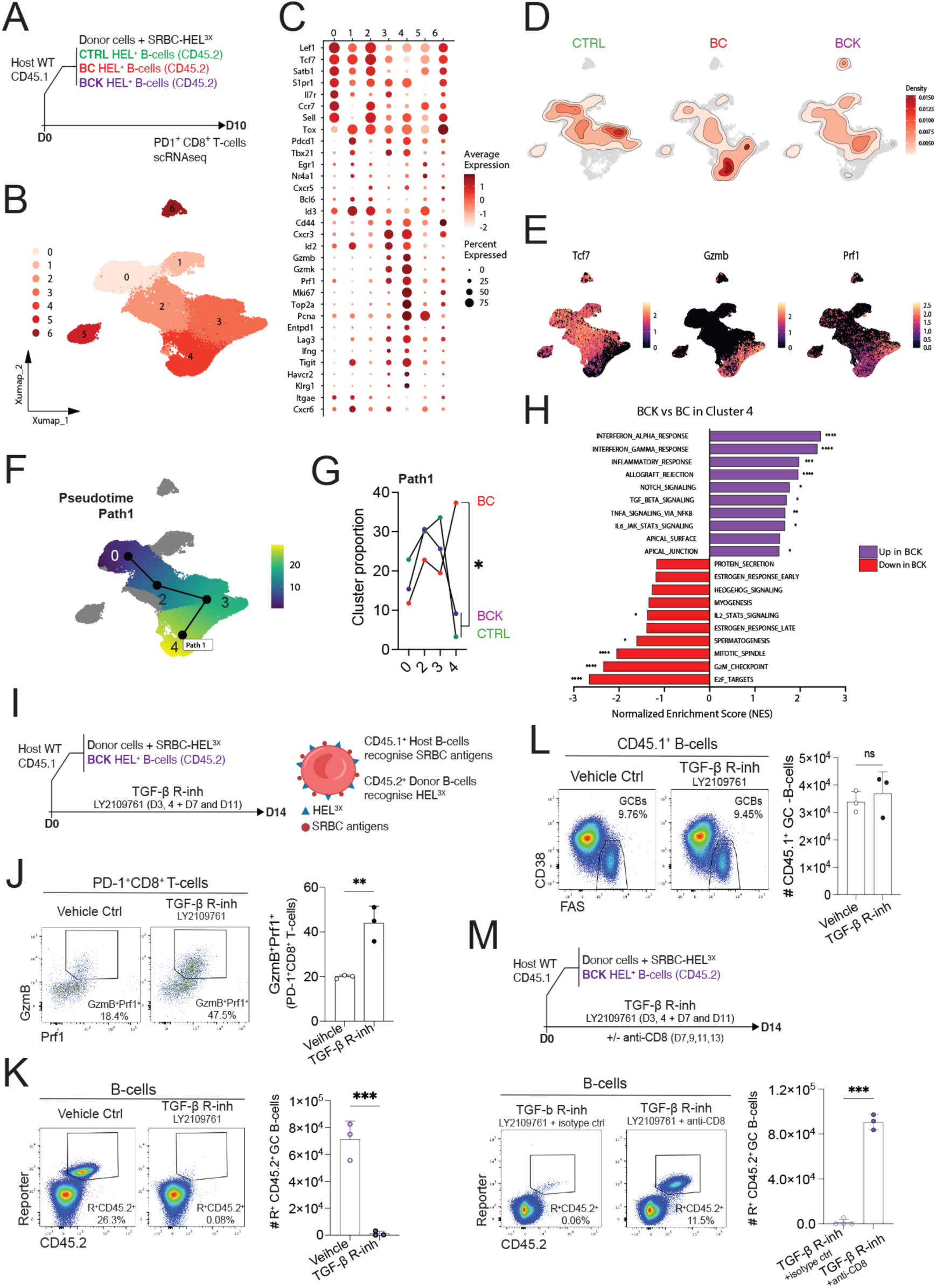
Immune-escape arises from impaired CD8⁺ T-cell effector differentiation. **(A)** Experimental design for single-cell RNA sequencing of PD-1⁺ CD8⁺ T-cells sorted from recipient mice 10 days after immunization. **(B)** UMAP projection of PD-1⁺ CD8⁺ T-cells colored by annotated subset identity. **(C)** Dot plot showing expression of selected marker genes across annotated CD8⁺ T-cell subsets. **(D)** Density plots showing the distribution of PD-1⁺CD8⁺ T-cells from Ctrl, BC, and BCK recipients across the UMAP. **(E)** UMAP projections showing expression of selected marker genes from (C). **(F)** Slingshot trajectory analysis of PD-1⁺ CD8⁺ T-cell differentiation, highlighting trajectory path1 from activated states toward stem-like, early effector and effector states. **(G)** Comparison of CD8⁺ T-cell subset distribution along path 1, which terminates in cytotoxic effector differentiation. Quantification shows mean cluster proportion from two biological replicates. **(H)** Gene set enrichment analysis of HALLMARK pathways comparing cluster 4 from BCK and BC conditions. **(I-K)** TGF-β receptor blockade following GC response derived from transferred BCK B-cells. **(I)** Experimental design for LY2109761 treatment. **(J)** Left, representative gating of granzymeB⁺ perforin⁺ effector cells among PD-1⁺ CD8⁺ T-cells. Right, quantification of PD-1⁺ CD8⁺ T-cells and granzymeB⁺ perforin⁺ effector CD8⁺ T-cells per 10⁶ splenocytes. **(K)** Left, representative gating of donor-derived reporter⁺ CD45.2⁺ B-cells. Right, quantification of donor-derived reporter⁺ CD45.2⁺ GC B-cells per 10⁶ splenocytes. **(L)** Endogenous CD45.1⁺ GC B-cell response to SRBCs in the absence or presence of TGF-β receptor inhibition with LY2109761. Left representative gating of host CD45.1⁺ GC B-cell B-cells. Right, quantification of host CD45.1⁺ GC B-cell B-cells. **(M)** Requirement for CD8⁺ T-cells during TGF-β receptor blockade. Top, schematic of experimental design. Bottom, representative gating of donor-derived reporter⁺ CD45.2⁺ B-cells from BCK recipients treated with LY2109761 together with isotype control or anti-CD8 antibody; and quantification of reporter⁺ CD45.2⁺ GC B-cells per 10⁶ splenocytes. Each symbol in **(J, L, K, M)** represents an individual mouse. Each symbol in **(G)** represents mean value from two biological replicates. **(J-L)** placebo n = 3, LY2109761 n = 3. **(M)** LY2109761 + isotype n = 3, LY2109761 + anti-CD8 n = 3. Small horizontal lines indicate mean ± SD. Data in **(J-L)** are representative of two independent experiments; data in **(K)** are representative of two independent experiments. *P ≤ 0.05, **P ≤ 0.01, ***P ≤ 0.001; Two-way ANOVA in **(G)**, value calculated on two biological replicates from each group, remaining statistical calculations used unpaired two-tailed Student’s t test. ns, not significant.

After integration of cells from all conditions, we identified seven transcriptionally distinct PD-1⁺ CD8⁺ T-cell clusters^53–57^ (**Fig.4B**). These clusters captured a continuum of activation and effector differentiation. Several clusters (0, 1, 2, and 6) retained high expression of stem-like or precursor-associated genes, including *Tcf7* and *Lef1*. Cluster 0 expressed *Il7r*, *Sell* and *Ccr7*, together with detectable *Tox* and *Id2*, consistent with an antigen-experienced stem-like state. Cluster 1 upregulated *Tox*, *Pdcd1*, *Tigit*, *Id2* and *Cxcr6*, with reduced *Il7r*, *Ccr7* and *Sell*, consistent with a progenitor-exhausted state. Cluster 2 retained a stem-like profile but expressed *Ccr7*, *Sell* and *Cxcr5*, suggesting lymphoid-homing capacity. Cluster 6 expressed Tcf7, *Lef1*, *Sell*, *Tox* and *Cd44*, consistent with a recently activated precursor-like PD-1⁺ CD8⁺ T-cell state (**Fig.4C-E**).

The remaining clusters (3, 4, 5) represented later stages of activation and effector differentiation. Cluster 3 expressed *Tbx21*, *Cd44* and *Id2* together with cytotoxic effector genes, including *Gzmb* and *Prf1*, consistent with an early effector precursor state. Cluster 5 retained stem-like features but showed elevated *Pcna* expression, indicative of a proliferative precursor-like population. By contrast, cluster 4 expressed *Havcr2* together with high levels of cytotoxic genes, including *Gzmb*, *Prf1* and *Ifng*, consistent with a terminal cytotoxic effector-like population (**Fig.4C-E**).

The distribution of PD-1⁺ CD8⁺ T-cell states differed markedly across conditions. Ctrl B-cell recipients were enriched for early activated and stem-like precursor populations, including clusters 0, 2, and 3. BC B-cell recipients showed a pronounced shift toward terminal cytotoxic differentiation, with marked accumulation of cluster 4. In contrast, BCK B-cell recipients preferentially accumulated progenitor-exhausted and precursor-like populations, including clusters 1, 6, 2, and 3, while showing reduced representation of terminal cytotoxic effectors (**Fig.4D**,**E**).

To resolve these differences more directly, we performed Slingshot trajectory analysis^58^. This revealed a differentiation continuum originating from cluster 0 and progressing through the stem-like cluster 2 before diverging into multiple fates (**Fig.S3A**). The dominant trajectory, Path 1, passed through the early effector state represented by cluster 3 and culminated in cluster 4, consistent with terminal cytotoxic differentiation (**Fig.S3A**,**4F**). Alternative trajectories terminated in the recently activated precursor-like cluster 6, the proliferative precursor-like cluster 5, or the progenitor-exhausted cluster 1 (**Fig.S3B-D**).

We next compared the distribution of cells along each inferred trajectory. The most prominent condition-dependent difference occurred along Path 1, the trajectory leading to terminal cytotoxic effector differentiation (**Fig.4G**,**S3B-D**). Both BC and BCK recipients showed progression from the antigen-experienced cluster 0 into the stem-like cluster 2 population, consistent with CD8⁺ T-cell activation. However, only BC recipients showed substantial progression to terminal cytotoxic effectors (cluster 4; **Fig.4G**). In BCK recipients, this transition was markedly reduced (**Fig.4G**). No comparable condition-dependent differences were observed along the alternative trajectories (**Fig.S3B-D**).

These data identify the immune-escape phenotype selected during premalignant GC B-cell evolution: CD8⁺ T-cells are recruited and activated but fail to complete terminal cytotoxic effector differentiation. Thus, evolved premalignant GC B-cells do not evade immune surveillance by preventing CD8⁺ T-cell entry or activation. Instead, they uncouple CD8⁺ T-cell activation from the acquisition of cytotoxic function required for premalignant GC B-cells elimination.

### TGF-β blockade restores CD8⁺ T-cell effector differentiation and surveillance

To identify pathways associated with impaired CD8⁺ T-cell effector differentiation in BCK recipients, we performed HALLMARK gene set enrichment analysis (GSEA) on scRNA-seq data from PD-1⁺ CD8⁺ T-cells. Because trajectory analysis showed that the major difference between BC and BCK recipients occurred along Path 1, the branch leading to terminal cytotoxic differentiation, we focused our analysis on terminal states within this effector trajectory (**Fig.4G**).

TGF-β signaling was among the pathways most prominently enriched in BCK-associated CD8⁺ T-cells (**Fig.4H**). TGF-β is well established to restrain CD8⁺ T-cell cytotoxic gene expression and effector function^59–61^, providing a plausible mechanistic basis for the failure of BCK-associated CD8⁺ T-cells to undergo full cytotoxic differentiation. Consistent with this interpretation, BCK-associated CD8⁺ T-cells showed reduced representation of metabolic and proliferative programs required to support effector maturation, including oxidative phosphorylation, fatty-acid metabolism, MYC targets, E2F targets and mTORC1 signaling (**Fig.4H**). Together, these data nominated TGF-β as a candidate suppressive pathway linking CD8⁺ T-cell activation to failed acquisition of a fully cytotoxic, surveillance-competent state.

We next asked whether TGF-β signaling was functionally responsible for the failure of CD8⁺ T-cells to eliminate BCK GC B-cells. To test this, we treated BCK-recipient mice with TGF-β receptor inhibitors during the GC response (**Fig.4I**,**S3E**). Treatment with LY2109761, a TGF-β receptor I/II kinase inhibitor, increased GrzmB⁺ Prf1⁺ PD-1⁺ CD8⁺ T-cell effectors compared with vehicle-treated controls (**Fig.4J**). Restoration of CD8⁺ T-cell effector differentiation was accompanied by a marked reduction in donor-derived BCK GC B-cells, which otherwise persisted at high levels in vehicle treated mice (**Fig.4K**). LY2109761 did not reduce endogenous CD45.1⁺ GC B-cells, arguing against non-specific GC toxicity and supporting selective elimination of the immune-evasive donor population (**Fig.4L**). Similar restoration of CD8⁺ T-cell effector differentiation was observed with vactosertib, an independent TGF-β type I receptor inhibitor, further supporting on-pathway activity (**Fig.S3E-H**).

Finally, to determine whether BCK GC B-cell elimination after TGF-β blockade was mediated by rescued CD8⁺ T-cell effector differentiation, we combined LY2109761 treatment with CD8⁺ T-cell depletion. In the absence of CD8⁺ T-cells, LY2109761 no longer reduced BCK GC B-cells (**Fig.4M**). Thus, TGF-β blockade removed the immune-evasive advantage of evolved premalignant GC B-cells by restoring CD8⁺ T-cell cytotoxic differentiation. These findings show that surveillance escape is reversible and reflects suppression of the CD8⁺ T-cell effector arm, rather than irreversible loss of immune recognition.

### Surveillance escape permits lymphoma-like GC B-cell evolution

BCK cells escaped CD8⁺ T-cell surveillance and persisted as expanded premalignant GC B-cell populations (**Fig.3**). We therefore asked whether this immune-evasive state created an opportunity for further malignant evolution. Because large-scale copy-number alterations are a hallmark of cancer^62^, we purified donor B-cells from Ctrl, BC, and BCK recipients and performed scRNA-seq (**Fig.5A**). We then inferred copy-number variation using inferCNV, with Ctrl donor B-cells as the reference population (**Fig.5B**).

**Figure 5.**
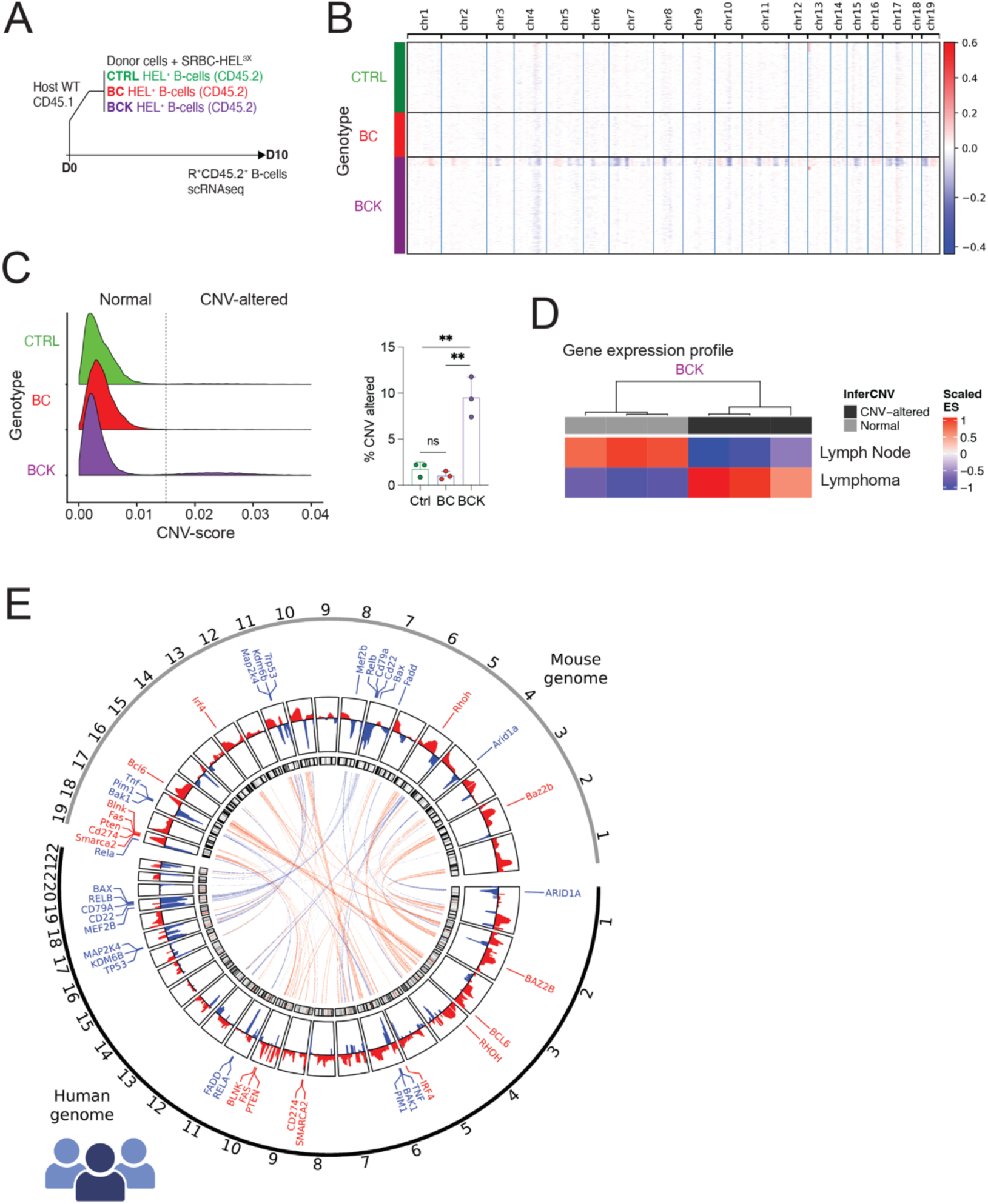
Surveillance escape permits lymphoma-like GC B-cell evolution. **(A)** Experimental design for single-cell RNA sequencing of reporter⁺ CD45.2⁺ donor B-cells sorted from Ctrl, BC, and BCK recipient mice 10 days after immunization. **(B)** Representative inferred copy-number variation analysis using scRNA-seq expression profiles from donor B-cells, with data integrated from three biological replicates per group. **(C)** Distribution of inferCNV scores across Ctrl, BC, and BCK conditions. Left, distribution of inferCNV scores across Ctrl, BC, and BCK conditions. An arbitrary threshold (indicated with dashed line) was applied to classify cells as normal or CNV-altered. Right, percentages of CNV-altered cells with high infer-CNV scores across Ctrl, BC, and BCK conditions. **(D)** Heatmap showing enrichment of normal lymph node and lymphoma gene signatures between normal and CNV-altered BCK cells. **(E)** Cross-species mapping of mouse inferCNV-predicted copy-number alterations onto syntenic regions of the human genome. Each symbol in **(C)** represents an individual mouse. Small horizontal lines indicate mean ± SD. **P ≤ 0.01; unpaired two-tailed Student’s t test.

Most BC and BCK B-cells lacked detectable large-scale CNV abnormalities (**Fig.5B**,**C**). However, a distinct subset, roughly 10%, of BCK B-cells were CNV-high, displaying broad chromosomal alterations (**Fig.5B**).

We next asked whether CNV-high BCK B-cells had also acquired lymphoma-like transcriptional features. To test this, we compared CNV-high and normal BCK B-cell populations with transcriptional signatures derived from normal lymph node B cells and B-cell lymphoma^63^. CNV-high BCK cells showed a clear enrichment toward the B-cell lymphoma signature, whereas normal BCK B-cells were closely aligned with normal lymph-node B cells (**Fig.5D**). Thus, lymphoma-like transcriptional identity was selectively associated with the CNV-high BCK B-cell subset, rather than with premalignant GC B-cell expansion alone.

To relate the mouse CNV patterns to human disease, we mapped predicted mouse alterations onto syntenic regions of the human genome. The altered regions corresponded to recurrent lymphoma-associated gains, including regions analogous to human 1q, 2p, 6p, 7q, 10q, and 18, and losses in regions analogous to 1p and 17p (**Fig.5E**). These regions overlap with copy-number alterations reported in human B-cell lymphoma^38,64–66^. Consistent with this convergence, several genes recurrently altered in B-cell lymphoma mapped to affected regions^67–69^, including predicted copy-number gain of BCL6 and predicted copy-number loss of TP53 and ARID1A.

Together, these findings indicate that escape from CD8⁺ T-cell surveillance allows fitness-enhanced premalignant GC B-cells to persist, creating an evolutionary window in which lymphoma-like genomic and transcriptional features can emerge. Thus, immune surveillance acts not only as a barrier to premalignant GC B-cell expansion, but also as a constraint on subsequent lymphoma-like evolution.

## Discussion

GCs are specialized lymphoid microenvironments in which B-cells undergo clonal expansion, somatic DNA mutation and affinity-based antibody selection^6–11^. These same processes create repeated opportunities for oncogenic lesions to arise^12–14^; nevertheless, lymphoma development is uncommon relative to the common detection of premalignant GC B-cells^15–18^. Our comparisons contrasting alterations associated with ISFN versus lymphoma suggests that this paradox reflects two linked selective pressures within GCs: acquisition of premalignant GC B-cell fitness and the need to survive local CD8⁺ T-cell surveillance.

In this framework, early ISFN-associated lesions generate premalignant GC B-cell states with increased fitness, allowing mutant B-cells to expand within GCs. However, these states are immune-surveilled and are eliminated by CD8⁺ T-cells. Further lymphoma-associated evolution converts this fitter-but-surveilled state into a fitter-and-surveillance-breaking state, in which CD8⁺ T-cells still infiltrate the premalignant GC and become activated, but fail to complete cytotoxic differentiation. Thus, premalignancy-to-lymphoma evolution selects not only for B-cells that grow better in the GC microenvironment, but for B-cells that can grow under local immune-pressure.

A central conclusion of this study is that GCs are neither immune-privileged nor immune-passive sites for lymphoma initiation. Although normal GCs contain few CD8⁺ T-cells, premalignant GC B-cell states triggered local CD8⁺ T-cell infiltration, activation and cytotoxic differentiation. In human ISFN, CD8⁺ T-cells were enriched in GCs containing BCL2⁺ premalignant B-cells, whereas NK-cells remained rare. In mouse models, premalignant GC B-cells similarly recruited only CD8⁺ T-cells into GCs and were selectively eliminated while physiological GC responses in the same mice were preserved. Thus, GCs can support a form of local immune-surveillance that discriminates premalignant from normal GC B-cells. This selectivity may explain how immune surveillance operates within a compartment where uncontrolled cytotoxicity would otherwise threaten antibody diversification and immunity.

This finding reframes follicular CD8⁺ T-cell biology in the context of lymphoma initiation. CXCR5⁺ follicular CD8⁺ T-cells have been described in chronic viral infection and advanced tumor contexts, where they can access B-cell follicles and contribute to the control of persistent viral reservoirs or cancer cells^55,70–74^. However, whether cytotoxic CD8⁺ T-cells function as a barrier to premalignant GC B-cell evolution has remained unclear. Our data show that CD8⁺ T-cells can enter premalignant GCs and restrain premalignant B-cell states before lymphoma development. The absence of NK-cell enrichment suggests that this is not a broad innate cytotoxic response, but rather an anatomically adapted CD8⁺ T-cell surveillance mechanism.

These findings place GC lymphomagenesis within the framework of cancer immunoediting. The classical immunoediting model proposes elimination, equilibrium and escape^75,76^. We found that premalignant GC B-cells initially entered an elimination-sensitive state: they expanded transiently but were removed by CD8⁺ T-cells. Human ISFN may represent a related contained or equilibrium-like state, in which premalignant B-cells persist within GCs but usually do not progress to lymphoma. Progression required a transition to immune escape, in which premalignant GC B-cells were no longer productively controlled. This is consistent with the view that the immune system not only eliminates cancer precursors but also shapes the evolutionary trajectories of emerging malignant clones^4^.

Immune escape at the premalignant GC stage differed from the dominant immune-evasion patterns described in established cancers. In established cancers, immune evasion is commonly associated with CD8⁺ T-cell dysfunction, exhaustion or spatial exclusion from tumor cells^29,77–79^, phenotypes that have been linked to TGF-β activity, including its capacity to remodel the stromal microenvironment and restrict productive T-cell trafficking^80–82^. By contrast, immune escape at the premalignant GC stage described here did not reflect absence of CD8⁺ T-cells, terminal exhaustion or spatial exclusion. CD8⁺ T-cells entered premalignant GCs and showed evidence of activation but failed to complete differentiation into cytotoxic effectors. Thus, in premalignant GCs, TGF-β appears to act primarily by restraining local cytotoxic licensing, whereas in established disease it may additionally contribute to broader tissue remodeling, stromal activation and immune-exclusion.

This context may be particularly relevant to lymphoma in later life. Lymphoma incidence increases markedly with age^83^. Ageing increases somatic mutation burden in normal and premalignant tissues, but mutation burden alone does not determine which premalignant cells progress^84^. Ageing also remodels secondary lymphoid organs, stromal signals and T-cell function, and elevated TGF-β signaling has been linked to age-dependent loss of CD8⁺ T-cell polyfunctionality^85,86^. Thus, lymphoma risk in later life may reflect two converging processes: increased generation/expansion of premalignant GC B-cells and reduced capacity of the aged lymphoid microenvironment to eliminate or contain these cells.

The finding that GCs are actively immune-surveilled also has important implications for how lymphoma-associated mutations and GC perturbation studies should be interpreted. Genetic lesions are often viewed through their effects on fitness, including survival, persistence, altered differentiation, GC retention or acquisition of additional genomic lesions. Our findings indicate that fitness within the GC also includes the ability to withstand or disable local immune surveillance. Conversely, loss of a genetically altered GC B-cell population need not always reflect reduced fitness or passive competitive failure; in some contexts, it may reflect increased immune visibility and active CD8⁺ T-cell-mediated elimination. GC B-cell genetics studies should therefore aim to distinguish B-cell-fitness from immune-surveillance sensitivity.

This work supports a framework for early immune interception. We do not propose that all ISFNs should be treated, because most cases do not progress and clinical risk stratification remains limited^15–18^. However, our data suggest that premalignant GCs should be evaluated not only by genetic composition but also by immune-surveillance status. A ISFN GC infiltrated by CD8⁺ T-cells that retain cytotoxic differentiation may remain under immune control, whereas an ISFN with CD8⁺ T-cell infiltration but impaired cytotoxic differentiation may represent a surveillance-breaking state. For individuals with ISFN, the greatest opportunity may lie before cancer becomes clinically advanced. Premalignant GC B-cells are likely to be more amenable to therapeutic control than established tumours, placing this disease window at the centre of cancer immune interception and precision prevention^3,87–89^. Understanding when and how premalignant GC B-cells shift from immune control to escape could identify individuals at risk of progression and create opportunities for earlier, less toxic, and more effective intervention.

## Methods

### Mouse strains

to model human ISFN GCs, we combined *BCL2*-overexpression with heterozygous *Crebbp*-loss. R26-BCL2^LSL^ mice^44^ were crossed with *Cd79a-Cre* and *Crebbp^fl/fl^* mice^43,46^ to generate B cells with conditional BCL2 overexpression alone (B), or BCL2 overexpression together with heterozygous Crebbp deletion (BC; *Crebbp^fl/+^*). To model a lymphoma-associated state, *Kmt2d^fl/fl^* mice^52^ were incorporated to generate *BCL2*-overexpressing, *Crebbp*-heterozygous, *Kmt2d*-deficient B-cells (BCK). These strains were further crossed to the SWHEL system to enable antigen-specific recruitment into germinal center (GC) responses following immunization with hen egg lysozyme (HEL) antigen^90^.

The following strains were obtained from The Jackson Laboratory (Bar Harbor, ME, USA): *C57BL/6J* (CD45.2, stock #000664), *B6.SJL-Ptprca Pepcb/BoyJ* (CD45.1, stock #002014), *Cd79a-Cre* (stock #020505), *Crebbp^fl/fl^* (stock #025178), and *Kmt2d^fl/fl^* (stock #032152). C57BL/6J and B6.SJL mice were maintained by in-house breeding at the Francis Crick Institute. *Cd79a-Cre* and *Rosa26^LSL-YFP^* mice were used as controls where indicated. Both male and female mice were included, with sex-matched cohorts in each experiment. All animals were housed under specific pathogen-free conditions at the Francis Crick Institute Biological Research Facility. All procedures were conducted in accordance with UK Home Office regulations and approved by the Francis Crick Institute Ethical Review Panel.

### Immunization, adoptive transfers, and *in vivo* treatments

for immunization in the SWHEL system, CD45.2^+^ HEL-binding donor B-cells were adoptively transferred into CD45.1⁺ congenic recipient mice. Briefly, 2×10⁵ HEL-binding B-cells were co-injected intravenously with2×10⁸ sheep red blood cells (SRBCs) conjugated to a recombinant HEL^3X^ protein. SRBCs in Alsever’s solution (TCS Bioscience) were conjugated to recombinant HEL^3X^ (R. Brink) using 1-ethyl-3-(3-dimethylaminopropyl) carbodiimide hydrochloride (Sigma), as described previously^91^. CD45.2⁺ splenic B-cells from SWHEL donor mice were purified by negative selection using anti-mouse CD43 (Ly-48) MicroBeads (Miltenyi Biotec). For flow cytometry and cell sorting after adoptive transfer, donor-derived CD45.2⁺ cells were enriched from recipient spleens by depleting CD45.1+ recipient cells using biotinylated anti-mouse CD45.1 antibody (clone A20, eBioscience), followed by Anti-Biotin MicroBeads (Miltenyi Biotec). For CD8+ T-cell depletion, mice received 100 μg anti-CD8 antibody (BioXCell, BE0061) or isotype control antibody (BioXCell, BE0090) by intraperitoneal injection in 200 μL PBS at the indicated time points. For TGF-β pathway inhibition, mice were treated with the dual TGF-β type I/II receptor inhibitor LY2109761 (MedChemExpress, HY-12075; 40 mg/kg). LY2109761 was dissolved in DMSO, diluted in 40% PEG300, 5% Tween-80 and 45% saline, and administered by oral gavage at the indicated time points. Vehicle-treated mice received the corresponding vehicle formulation. In independent experiments, mice were treated with vactosertib (MedChemExpress, HY-19928; 2.5 mg/kg), dissolved in DMSO and diluted in 90% corn oil, and administered by oral gavage at the indicated time points, with matched vehicle controls.

### Flow cytometry and cell sorting

single-cell suspensions of spleen were prepared in FACS buffer (2% FBS, 2 mM EDTA, in PBS; Gibco) and were treated with ACK buffer (Gibco) for erythrocyte lysis. Single-cell suspensions were stained with antibodies. The use of biotinylated antibodies was followed by incubation with fluorochrome-conjugated streptavidin (1:200 dilution). For the analyses of SWHEL mice, HEL-binding cells were stained with 50 ng/mL HEL (Sigma) or HEL^3X^ (R. Brink) together with anti-mouse IgG1 antibody as the first step, followed by blocking with normal mouse serum (Sigma) before incubation with anti-HEL antibody HyHEL9-AF647 (R. Brink). Dead cells were excluded using Zombie NIR™ Fixable Viability Kit (BioLegend). Samples were acquired on an LSR-Fortessa (BD Biosciences) with FACS-Diva software (BD Biosciences) and data were analyzed with FlowJo software (v10.8.1, Tree Star). Red blood cell-lysed splenocytes (1 × 10⁷ cells) were resuspended in 1 mL complete culture medium (DMEM supplemented with 10% FCS, non-essential amino acids, sodium pyruvate, HEPES, and β-mercaptoethanol). Cells were stimulated for 1-1.5 hours in the presence of Cell Stimulation Cocktail (plus protein transport inhibitors; ThermoFisher, 00-4970-03). Following stimulation, cells were pelleted by centrifugation and stained with a surface antibody cocktail in the presence of Fc block. Cells were then fixed, permeabilized, and subjected to intracellular staining for granzymes and perforin. For cell sorting, single-cell suspensions were prepared from spleens after red blood cell lysis and stained for surface markers. Reporter⁺ donor B-cells and PD-1⁺CD8⁺ T-cells were sorted from recipient mice 10 days after immunization with SRBCs-HEL^3X^. Where indicated, both populations were sorted from the same recipient mouse, mixed at equal numbers and loaded onto the same 10x Genomics run. B-cell and T-cell populations were subsequently separated computationally using canonical lineage markers and immune-receptor transcript content.

### Multiplex immunofluorescence

for human samples, control lymph node (LN, n = 7) and ISFN (n = 6) tissue sections were prepared from FFPE blocks provided by the Imperial College Healthcare Tissue Bank (Project No. R24018). FFPE tissue sections (3 μm) were baked for 1h and processed on the Leica Bond RX platform. Endogenous peroxidase activity was blocked using 3% hydrogen peroxide, followed by protein blocking with 0.1% BSA in PBST. Antigen retrieval and antibody stripping between sequential staining cycles were performed using Epitope Retrieval Solution 2 (pH 6; Leica, AR9961) for 20 minutes at 95°C. Multiplex immunofluorescence (6-plex) staining was conducted using Opal fluorophores with the following antibody–fluorophore pairings and staining order: NCAM-1 (Abcam, ab237708; 1:2000) with Opal 520 (1:300), BCL6 (Cell Signaling Technology, 5650S; 1:100) with Opal 570 (1:300), CD10 (Abcam, ab256494; 1:200) with Opal 620 (1:300), CD8 (Dako, GA623; ready-to-use) with Opal 480 (1:200), BCL2 (Abcam, ab182858; 1:500) with Opal 690 (1:150), and CD20 (Abcam, ab9475; 1:200) with Opal 780 (1:100). Secondary detection was performed using the Novolink Polymer Detection System (Leica, RE7260-CE, ready-to-use), compatible with both mouse- and rabbit-derived primary antibodies. Slides were counterstained with DAPI (Thermo Scientific, 62248; 1:2500) and mounted using ProLong Gold Antifade reagent (Invitrogen, P36934). Whole-slide imaging was performed using the PhenoImager HT (formerly Vectra Polaris) automated imaging system with PhenoImager HT 2.1 software. Spectral unmixing was performed using the integrated software feature, incorporating a tissue-specific autofluorescence control slide for background subtraction. For mouse tissues, spleens were fixed in 4% paraformaldehyde (PFA) for 24 hours, transferred to 70% ethanol, and embedded in paraffin for FFPE block preparation. FFPE tissue sections (3 μm) were processed as described for human samples. Multiplex immunofluorescence (6-plex) was performed, and antibodies were applied with Opal™ pairings in the following order: 1:300, BCL6 (Cell Signaling Technology, 5650S) at 1:100 with Opal 480 at 1:200, GFP (Thermo Fisher Scientific, A-6455) at 1:500 with Opal 520 at 1:300, PD-1 (Abcam, ab214421) at 1:500 with Opal 620 at 1:300, and CD8 (Abcam, ab217344) at 1:500 with Opal 690 at 1:150. All exported unmixed images were analyzed using QuPath (v0.6.0-arm64). To analyze cells within germinal centers (GCs), BCL6^+^ regions were first annotated as areas of interest. Cells within these regions were segmented using the InstanSeg extension, incorporating all staining channels from the slides. CD8^+^, BCL2^+^, GFP^+^ and cells were quantified within each annotated GC and reported either as total cells per GC or normalized to GC area, as indicated in the corresponding figure legends.

### Single cell RNA sequencing

reporter⁺ donor-derived B-cells and PD-1⁺CD8⁺ T-cells were sorted using a FACSAria II cell sorter (BD Immunocytometry Systems) ) from recipient mice 10 days after immunization. For each experimental group (Ctrl, BC and BCK), two biological replicates were included. Where both B- and T-cells were profiled from the same mouse, equal numbers of sorted reporter^+^ B-cells and PD-1^+^CD8^+^ T cells were combined before loading onto 10x Genomics Chromium chips. Approximately 10,000 cells from each population were targeted per sample. In some cases, samples were split across two 10x lanes/chips to increase total cell recovery. Library preparation and sequencing were performed by the Francis Crick Institute according to the manufacturer’s instructions for the Chromium Next GEM Single Cell 5′ workflow (v2 or v3, as applicable). Gene-expression libraries and, where applicable, immune-receptor V(D)J libraries were generated. Libraries were sequenced on an Illumina NovaSeq 6000 platform using S2 flow cells where applicable.

### scRNAseq processing of T and B lymphocytes

FASTQ files were aligned to the refdata-gex-mm10-2020-A mouse reference genome using the 10X Genomics CellRanger pipeline (v7.0.1). Processing of scRNAseq data was performed following the rapids_singlecell (v0.11.1) workflow in Python (v3.12.12). For quality control, the gene expression matrices from Ctrl, BC, and BCK mice were assessed separately to flag and remove low-quality cells. This includes empty droplets (n_genes_by_counts < 1000), doublets (n_genes_by_counts > 6000, total_counts > 30000, dead cells (pct_counts_MT > 10%) and contaminant erythrocytes (pct_counts_HG > 0.1%). To ensure data quality, doublets identified by scDblFinder (v1.14.0) in R (v4.3.2), and cells co-expressing > 0.1% of TCR reads and > 5% BCR reads, were further filtered out. RNA decontamination using DecontX under the celda package (v1.16.1), where contaminated cells (DecontX contamination score > 0.2) were removed from each sample. TCR and BCR reads were excluded from gene expression to prevent clustering by VDJ clonotypes.

Decontaminated gene expression matrices were merged and log1p normalised. scVI from scvi-tools (v1.4.1) integration was performed with regression on the following covariates: S_score, G2M_score, pct_counts_MT and pct_counts_RB to segregate cells by immune cell type identity rather than proliferation and metabolic states. Using the latent space from scVI, Leiden clusters were identified with a resolution of 0.4 and were annotated as T cells (Cd3d, Cd3e, Cd8a); B-cells (Cd19, Ms4a1, Cd79a); and plasma cells (Sdc1, Jchain, Prdm1); and dendritic cells (Itgax, Flt3, Zbtb46). T-cell clusters were isolated and further re-clustered to identify distinct T-cell subpopulations following an identical dimensionality and data integration approach without any covariate regressions, using a resolution of 0.2. T-cell subsets were annotated according to genes shown in Figure 4C.

### Functional characterization of T-cell lineages between BC and BCK

Differential gene expression between BCK and BC was calculated for all divergent T-cell subpopulation from each lineage with the ‘FindMarkers()’ function from Seurat using a zero-inflated generalized linear model (MAST, v1.26.0) method. All differentially expressed genes were pre-ranked based on average log2 fold change expression. Gene set enrichment analysis was performed using the ‘GSEA()’ function from the clusterProfiler package (v4.8.2) on the MSigDB HALLMARK gene sets, with parameters: nperm = 1000, min.overlap = 10. Pathways are considered significant when adjusted q-value < 0.05.

### Trajectory inference of T-cells

Trajectory analysis of T-cells was performed using slingshot packages (v2.8.0) on the T-cell UMAP embeddings, with cluster 0 specified as the starting cluster. Lineages were inferred with ‘getLineages()’ and ‘getCurves()’ function with default parameters. Inferred trajectories were visualized by overlaying the minimum spanning tree on the UMAP embeddings.

### Copy number variation inference of BC and BCK B-cells

copy number variations (CNVs) in B-cells from BC and BCK mice were analyzed using the infercnvpy algorithm (v0.6.1) with window_size set to 158 (1% of unique genes captured), step=10 and calculate_gene_values = True. Reporter+ Ctrl donor B-cells were used as the reference population to establish baseline expression. CNV score for each cell was calculated as the average CNV value per gene, with a threshold set to distinguish normal cells and CNV-altered cells (cnv_score > 0.015). MGI symbols were converted to HGNC orthologs, including one-to-many ortholog mapping, using BioMart (v2.58.0). Mouse CNV regions were projected onto the human reference genome (GRCh38/hg38, annotated with GENCODE v27) to assess conserved syntenic relationships between mouse and human loci.

### Cross-species FL signature scoring

Normal lymph node B-cell (LN) and FL samples were retrieved from the Gómez-Abad et al (GSE32018) microarray dataset and reanalysed using limma (v3.56.2). To derive LN and FL transcriptional signatures, differentially expressed genes were selected using thresholds of Benjamini–Hochberg (BH) adjusted P < 0.05, log2 fold-change > 0.5, with a mean expression > 0.5 in LN or FL samples. The top 200 genes for both LN and FL were ranked by adjusted P value (Table S1). To calculate the FL enrichment score, the mouse single-cell transcriptomes were pseudo-bulked by infercnv clusters and individual BCK mice using the ‘AggregateExpression()’ from Seurat. After orthologue gene conversion across species, enrichment of the human FL signature in mouse pseudo-bulk profiles was quantified with ssGSEA using the GSVA package (v1.51.6).

### Statistical analysis

Statistical methods used for each comparison are specified in the corresponding figure legends. Statistical analyses were performed using GraphPad Prism 11 or R using rstatix (v0.7.2) and ggpubr (v0.6.3). Unless otherwise stated, tests were two-sided and p values or adjusted p values < 0.05 were considered statistically significant. Biological replicate numbers are indicated in the figure legends. Experiments were performed with at least three biological replicates unless otherwise stated. For the PD-1+CD8+ T-cell scRNA-seq analysis, the replicate-2 T-cell libraries from all three experimental groups were excluded because of technical failure during FACS sorting; matched B-cell libraries were retained unless otherwise stated.

## Acknowledgements

We thank the members of the Immunity and Cancer laboratory [Francis Crick Institute (FCI), London, UK] for critical discussions and comments. We thank Arianne Richard and Martin Turner [Babraham Institute] for critical discussions and comments. We thank the FCI scientific platforms (Biological Resource Facility, Flow Cytometry, Histopathology) for expert advice and technical support. We acknowledge the Genomics Science Technology Platform at The Francis Crick Institute, for their contribution to the single-cell capture experiment, library construction, and sequencing, particularly Hubert Slawinski, Deb Jackson, Sam Jones and Marg Crawford. Schematics were created with BioRender.com

## Author contributions

Conceptualization: L.Z., and D.P.C.

Methodology: L.Z., M.S.H.

Investigation: L.Z., M.S.H., O. A., P. A., A. S., V. B.

Resources: H.K., H.C.R., J.O., D.P.C.

Visualization: L.Z., M.S.H.

Funding acquisition: O.A., H.C.R., J.F., J.O., D.P.C.

Supervision: D.P.C.

Writing-original draft: L.Z., M.S.H., and D.P.C.

Writing-review and editing: All authors.

---

**Figure S1.**
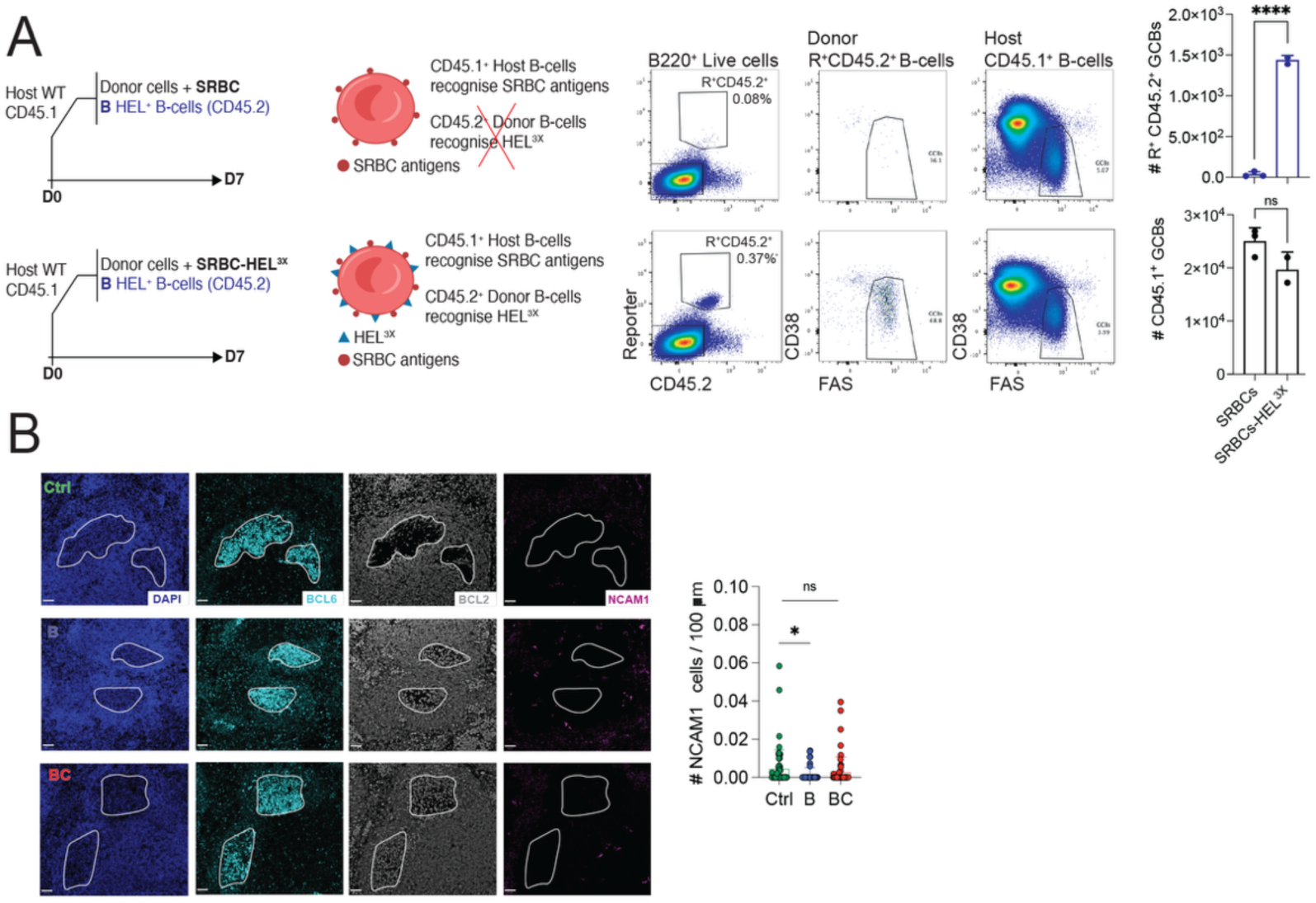
Donor- and host-derived GC B-cell responses. **(A)** Antigen-specific recruitment of donor-derived SWHEL B-cells into GC responses. Top: left, schematic of experiment designed to test whether HEL^+^ B-cells respond to SRBC antigens; right, representative gating and quantification of reporter⁺ CD45.2⁺ donor SWHEL B-cells and derived GC B-cells. Bottom: left, schematic of experiment design to test whether HEL^+^ B-cells respond to SRBC-HEL^3x^ immunization; right, representative gating and quantification of reporter⁺ CD45.2⁺ donor SWHEL B-cells and derived GC B cells. **(B)** Left, immunofluorescence analysis of spleen sections from immunized recipient mice 7 days after transfer of Ctrl, B, or BC SWHEL B-cells. Representative counterstains showing DAPI, BCL6⁺, BCL2^+^, NCAM1⁺ cells (DAPI: blue; BCL6, cyan; BCL2, white; NCAM1, purple). Scale bar, 50 μm. Right, quantification of NCAM1⁺ cells per GC. Each symbol in **(A)** represents an individual mouse n = 2 to 3 mice. **(B)** are representative images from 3 mice. Small horizontal lines indicate mean ± SD.

**Figure S2.**
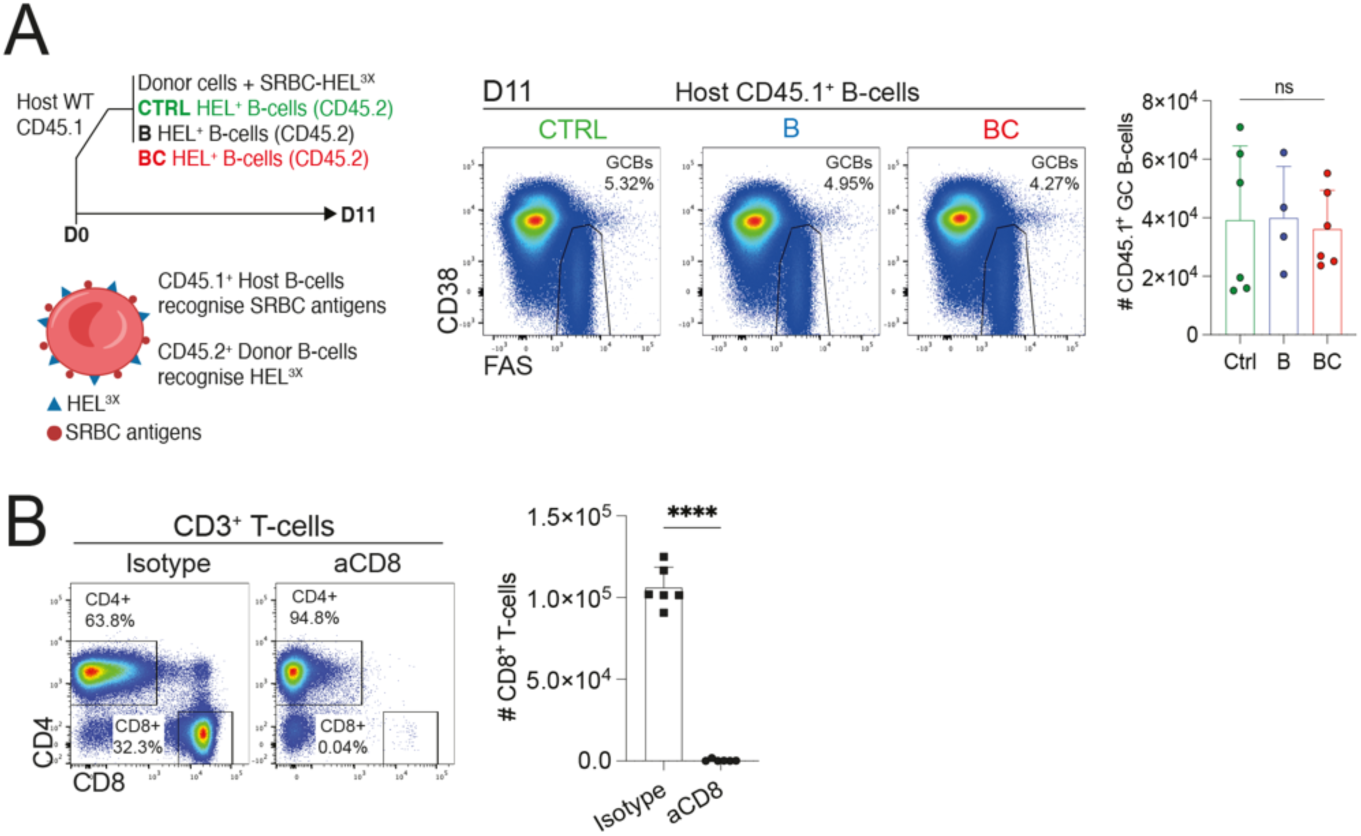
Host-derived GC B-cells are not eliminated by CD8⁺ T-cells. **(A)** Endogenous CD45.1⁺ GC B-cell responses 11 days after immunization in recipients of Ctrl, B, or BC donor B-cells. Left, schematic of experimental design. Right, representative gating of endogenous CD45.1⁺ GC B-cells and quantification of endogenous GC B-cell numbers per 10⁶ splenocytes. **(B)** Specificity of CD8⁺ T-cell depletion 14 days after immunization. Left, representative gating of CD8⁺ T-cells from recipient splenocytes treated with isotype control or anti-CD8 antibody. Right, quantification of CD8⁺ T-cell numbers per 10⁶ splenocytes. Each symbol in **(A and B)** represents an individual mouse. **(A)** Ctrl n = 6, B n = 4, BC n = 6. **(B)** isotype n = 6, anti-CD8 n = 6. Small horizontal lines indicate mean ± SD. Data in **(A)** are from two independent experiments; data in **(B)** are representative of two independent experiments. ****P ≤ 0.0001; unpaired two-tailed Student’s t test. ns, not significant.

**Figure S3.**
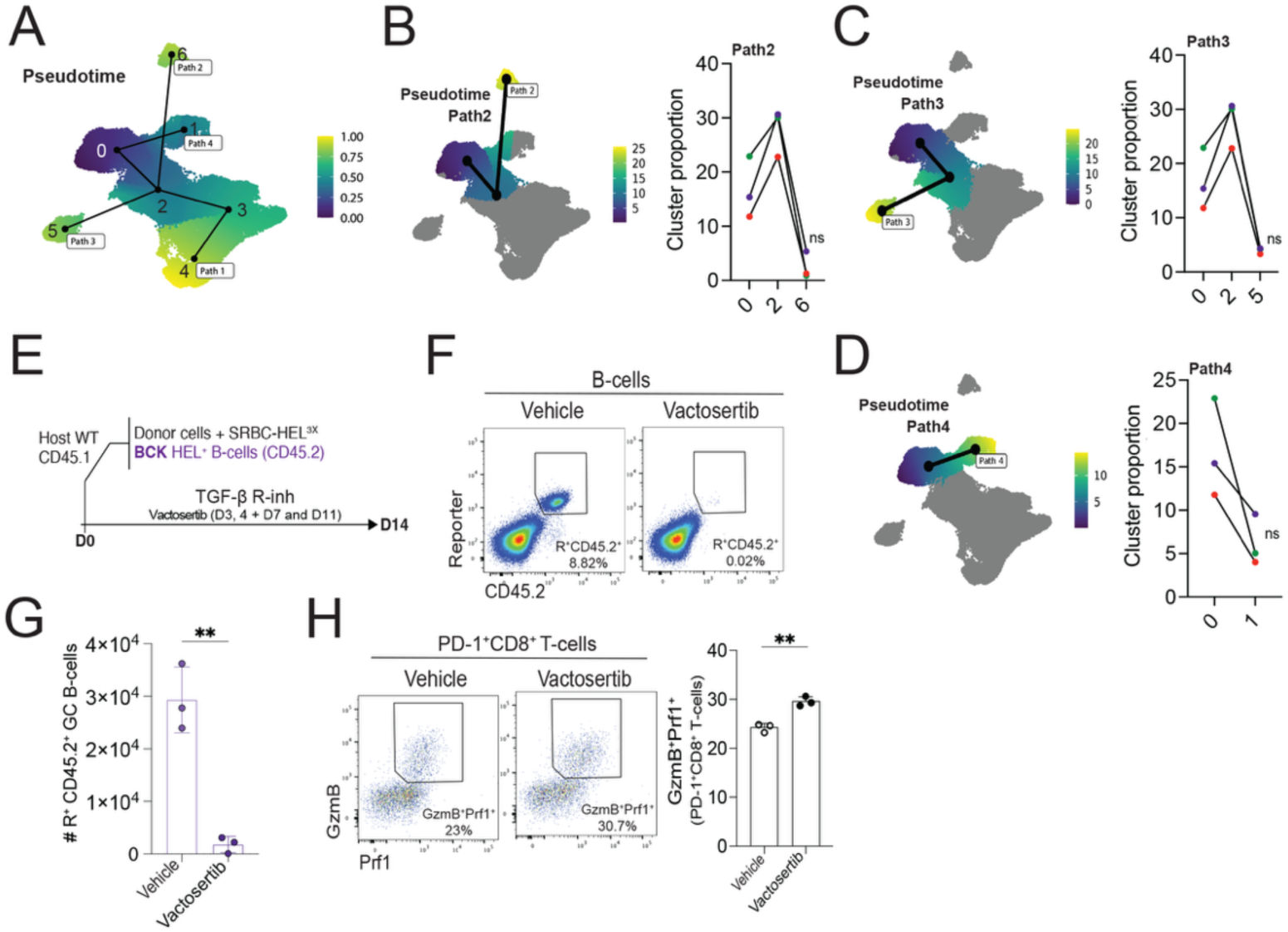
TGF-β blockade restores CD8⁺ T-cell effector differentiation. **(A)** Slingshot trajectory analysis of PD-1⁺ CD8⁺ T-cell differentiation, showing all retrieved trajectories from slingshot analysis. **(B)** Slingshot trajectory analysis of PD-1⁺CD8⁺ T-cell differentiation, highlighting trajectory path2 that terminates at the recently activated precursor-like cells (left) and comparison of CD8⁺ T-cell subset distribution along this path (right). **(C)** Slingshot trajectory analysis of PD-1⁺CD8⁺ T-cell differentiation, highlighting trajectory path3 that terminates at proliferative precursor-like cells (left) and comparison of CD8⁺ T-cell subset distribution along this path (right). **(D)** Slingshot trajectory analysis of PD-1⁺CD8⁺ T-cell differentiation, highlighting trajectory path4 that terminates at the progenitor-exhausted cells (left) and comparison of CD8⁺ T-cell subset distribution along this path (right). **(E-H)** TGF-β receptor blockade following GC response derived from transferred BCK B-cells. **(E)** Experimental design for Vactosertib treatment. **(F)** Representative gating of donor-derived reporter⁺ CD45.2⁺ B-cells. **(G)** Quantification of donor-derived reporter⁺ CD45.2⁺ GC B-cells per 10⁶ splenocytes. **(H)** Left, representative gating of granzymeB⁺ perforin⁺ effector cells among PD-1⁺ CD8⁺ T-cells. Right, quantification of PD-1⁺ CD8⁺ T-cells and granzymeB⁺ perforin⁺ effector CD8⁺ T-cells per 10⁶ splenocytes. Each symbol in **(G, H)** represents an individual mouse. Each symbol in **(B-D)** represents mean value from two biological replicates. **(G, H)** placebo n = 3, vactosertib n = 3. Small horizontal lines indicate mean ± SD. Data in **(G-H)** is from one experiment. **P ≤ 0.01; Two-way ANOVA in **(G)**, value calculated on two biological replicates from each group, other analysis: unpaired two-tailed Student’s t test. ns, not significant.

## References

1 Greaves, M. & Maley, C. C. Clonal evolution in cancer. Nature 481, 306–313 (2012). 10.1038/nature10762

2 Curtius, K., Wright, N. A. & Graham, T. A. Evolution of Premalignant Disease. Cold Spring Harb Perspect Med 7 (2017). 10.1101/cshperspect.a026542

3 Rane, J. K., Frankell, A. M., Weeden, C. E. & Swanton, C. Clonal Evolution in Healthy and Premalignant Tissues: Implications for Early Cancer Interception Strategies. Cancer Prev Res (Phila) 16, 369–378 (2023). 10.1158/1940-6207.CAPR-22-0469

4 Gubin, M. M. & Vesely, M. D. Cancer Immunoediting in the Era of Immuno-oncology. Clin Cancer Res 28, 3917–3928 (2022). 10.1158/1078-0432.CCR-21-1804

5 de Visser, K. E. & Joyce, J. A. The evolving tumor microenvironment: From cancer initiation to metastatic outgrowth. Cancer Cell 41, 374–403 (2023). 10.1016/j.ccell.2023.02.016

6 Weisel, F. J., Zuccarino-Catania, G. V., Chikina, M. & Shlomchik, M. J. A Temporal Switch in the Germinal Center Determines Differential Output of Memory B and Plasma Cells. Immunity 44, 116–130 (2016). 10.1016/j.immuni.2015.12.004

7 Schiepers, A. et al. Molecular fate-mapping of serum antibody responses to repeat immunization. Nature 615, 482–489 (2023). 10.1038/s41586-023-05715-3

8 Ise, W. & Kurosaki, T. Plasma cell differentiation during the germinal center reaction. Immunol Rev 288, 64–74 (2019). 10.1111/imr.12751

9 Mesin, L. et al. Restricted Clonality and Limited Germinal Center Reentry Characterize Memory B Cell Reactivation by Boosting. Cell 180, 92–106 e111 (2020). 10.1016/j.cell.2019.11.032

10 Victora, G. D. & Nussenzweig, M. C. Germinal Centers. Annu Rev Immunol 40, 413–442 (2022). 10.1146/annurev-immunol-120419-022408

11 Nakagawa, R. & Calado, D. P. Positive Selection in the Light Zone of Germinal Centers. Front Immunol 12, 661678 (2021). 10.3389/fimmu.2021.661678

12 Sungalee, S. et al. Germinal center reentries of BCL2-overexpressing B cells drive follicular lymphoma progression. J Clin Invest 124, 5337–5351 (2014). 10.1172/JCI72415

13 Dominguez, P. M. et al. TET2 Deficiency Causes Germinal Center Hyperplasia, Impairs Plasma Cell Differentiation, and Promotes B-cell Lymphomagenesis. Cancer Discov 8, 1632–1653 (2018). 10.1158/2159-8290.CD-18-0657

14 Mlynarczyk, C., Fontan, L. & Melnick, A. Germinal center-derived lymphomas: The darkest side of humoral immunity. Immunol Rev 288, 214–239 (2019). 10.1111/imr.12755

15 Henopp, T., Quintanilla-Martinez, L., Fend, F. & Adam, P. Prevalence of follicular lymphoma in situ in consecutively analysed reactive lymph nodes. Histopathology 59, 139–142 (2011). 10.1111/j.1365-2559.2011.03897.x

16 Oishi, N., Segawa, T., Miyake, K., Mochizuki, K. & Kondo, T. Incidence, clinicopathological features and genetics of in-situ follicular neoplasia: a comprehensive screening study in a Japanese cohort. Histopathology 80, 820–826 (2022). 10.1111/his.14617

17 Bermudez, G. et al. Incidental and Isolated Follicular Lymphoma In Situ and Mantle Cell Lymphoma In Situ Lack Clinical Significance. Am J Surg Pathol 40, 943–949 (2016). 10.1097/PAS.0000000000000628

18 Jegalian, A. G. et al. Follicular lymphoma in situ: clinical implications and comparisons with partial involvement by follicular lymphoma. Blood 118, 2976–2984 (2011). 10.1182/blood-2011-05-355255

19 Swann, J. B. & Smyth, M. J. Immune surveillance of tumors. J Clin Invest 117, 1137–1146 (2007). 10.1172/JCI31405

20 Corthay, A. Does the immune system naturally protect against cancer? Front Immunol 5, 197 (2014). 10.3389/fimmu.2014.00197

21 Smyth, M. J. et al. Perforin-mediated cytotoxicity is critical for surveillance of spontaneous lymphoma. J Exp Med 192, 755–760 (2000). 10.1084/jem.192.5.755

22 Street, S. E., Trapani, J. A., MacGregor, D. & Smyth, M. J. Suppression of lymphoma and epithelial malignancies effected by interferon gamma. J Exp Med 196, 129–134 (2002). 10.1084/jem.20020063

23 Bax, H. I. et al. B-cell lymphoma in a patient with complete interferon gamma receptor 1 deficiency. J Clin Immunol 33, 1062–1066 (2013). 10.1007/s10875-013-9907-0

24 Brennan, A. J., Chia, J., Trapani, J. A. & Voskoboinik, I. Perforin deficiency and susceptibility to cancer. Cell Death Differ 17, 607–615 (2010). 10.1038/cdd.2009.212

25 Miles, B. et al. Follicular Regulatory CD8 T Cells Impair the Germinal Center Response in SIV and Ex Vivo HIV Infection. PLoS Pathog 12, e1005924 (2016). 10.1371/journal.ppat.1005924

26 Fukazawa, Y. et al. B cell follicle sanctuary permits persistent productive simian immunodeficiency virus infection in elite controllers. Nat Med 21, 132–139 (2015). 10.1038/nm.3781

27 Connick, E. et al. Compartmentalization of simian immunodeficiency virus replication within secondary lymphoid tissues of rhesus macaques is linked to disease stage and inversely related to localization of virus-specific CTL. J Immunol 193, 5613–5625 (2014). 10.4049/jimmunol.1401161

28 Connick, E. et al. CTL fail to accumulate at sites of HIV-1 replication in lymphoid tissue. J Immunol 178, 6975–6983 (2007). 10.4049/jimmunol.178.11.6975

29 Laurent, C., Dietrich, S. & Tarte, K. Cell cross talk within the lymphoma tumor microenvironment: follicular lymphoma as a paradigm. Blood 143, 1080–1090 (2024). 10.1182/blood.2023021000

30 Ame-Thomas, P. et al. CD10 delineates a subset of human IL-4 producing follicular helper T cells involved in the survival of follicular lymphoma B cells. Blood 125, 2381–2385 (2015). 10.1182/blood-2015-02-625152

31 Ame-Thomas, P. et al. Characterization of intratumoral follicular helper T cells in follicular lymphoma: role in the survival of malignant B cells. Leukemia 26, 1053–1063 (2012). 10.1038/leu.2011.301

32 Mourcin, F. et al. Follicular lymphoma triggers phenotypic and functional remodeling of the human lymphoid stromal cell landscape. Immunity 54, 1788–1806 e1787 (2021). 10.1016/j.immuni.2021.05.019

33 Townsend, W. et al. The architecture of neoplastic follicles in follicular lymphoma; analysis of the relationship between the tumor and follicular helper T cells. Haematologica 105, 1593–1603 (2020). 10.3324/haematol.2019.220160

34 Pillai, R. K., Surti, U. & Swerdlow, S. H. Follicular lymphoma-like B cells of uncertain significance (in situ follicular lymphoma) may infrequently progress, but precedes follicular lymphoma, is associated with other overt lymphomas and mimics follicular lymphoma in flow cytometric studies. Haematologica 98, 1571–1580 (2013). 10.3324/haematol.2013.085506

35 Pasqualucci, L. et al. Inactivating mutations of acetyltransferase genes in B-cell lymphoma. Nature 471, 189–195 (2011). 10.1038/nature09730

36 Schmidt, J. et al. CREBBP gene mutations are frequently detected in in situ follicular neoplasia. Blood 132, 2687–2690 (2018). 10.1182/blood-2018-03-837039

37 Vogelsberg, A. et al. Genetic evolution of in situ follicular neoplasia to aggressive B-cell lymphoma of germinal center subtype. Haematologica 106, 2673–2681 (2021). 10.3324/haematol.2020.254854

38 Bai, B. et al. Multi-omics profiling of longitudinal samples reveals early genomic changes in follicular lymphoma. Blood Cancer J 14, 147 (2024). 10.1038/s41408-024-01124-5

39 Pasqualucci, L. et al. Genetics of follicular lymphoma transformation. Cell Rep 6, 130–140 (2014). 10.1016/j.celrep.2013.12.027

40 Okosun, J. et al. Integrated genomic analysis identifies recurrent mutations and evolution patterns driving the initiation and progression of follicular lymphoma. Nat Genet 46, 176–181 (2014). 10.1038/ng.2856

41 Montes-Moreno, S. et al. Intrafollicular neoplasia/in situ follicular lymphoma: review of a series of 13 cases. Histopathology 56, 658–662 (2010). 10.1111/j.1365-2559.2010.03529.x

42 Garcia-Ramirez, I. et al. Crebbp loss cooperates with Bcl2 overexpression to promote lymphoma in mice. Blood 129, 2645–2656 (2017). 10.1182/blood-2016-08-733469

43 Hobeika, E. et al. Testing gene function early in the B cell lineage in mb1-cre mice. Proc Natl Acad Sci U S A 103, 13789–13794 (2006). 10.1073/pnas.0605944103

44 Knittel, G. et al. B-cell-specific conditional expression of Myd88p.L252P leads to the development of diffuse large B-cell lymphoma in mice. Blood 127, 2732–2741 (2016). 10.1182/blood-2015-11-684183

45 Kuppers, R. & Dalla-Favera, R. Mechanisms of chromosomal translocations in B cell lymphomas. Oncogene 20, 5580–5594 (2001). 10.1038/sj.onc.1204640

46 Kang-Decker, N. et al. Loss of CBP causes T cell lymphomagenesis in synergy with p27Kip1 insufficiency. Cancer Cell 5, 177–189 (2004). 10.1016/s1535-6108(04)00022-4

47 Schroers-Martin, J. G. et al. Tracing Founder Mutations in Circulating and Tissue-Resident Follicular Lymphoma Precursors. Cancer Discov 13, 1310–1323 (2023). 10.1158/2159-8290.CD-23-0111

48 Srinivas, S. et al. Cre reporter strains produced by targeted insertion of EYFP and ECFP into the ROSA26 locus. BMC Dev Biol 1, 4 (2001). 10.1186/1471-213x-1-4

49 Phan, T. G. et al. B cell receptor-independent stimuli trigger immunoglobulin (Ig) class switch recombination and production of IgG autoantibodies by anergic self-reactive B cells. J Exp Med 197, 845–860 (2003). 10.1084/jem.20022144

50 Schaerli, P. et al. CXC chemokine receptor 5 expression defines follicular homing T cells with B cell helper function. J Exp Med 192, 1553–1562 (2000). 10.1084/jem.192.11.1553

51 Breitfeld, D. et al. Follicular B helper T cells express CXC chemokine receptor 5, localize to B cell follicles, and support immunoglobulin production. J Exp Med 192, 1545–1552 (2000). 10.1084/jem.192.11.1545

52 Lee, J. E. et al. H3K4 mono- and di-methyltransferase MLL4 is required for enhancer activation during cell differentiation. Elife 2, e01503 (2013). 10.7554/eLife.01503

53 Siddiqui, I. et al. Intratumoral Tcf1(+)PD-1(+)CD8(+) T Cells with Stem-like Properties Promote Tumor Control in Response to Vaccination and Checkpoint Blockade Immunotherapy. Immunity 50, 195–211 e110 (2019). 10.1016/j.immuni.2018.12.021

54 Utzschneider, D. T. et al. T Cell Factor 1-Expressing Memory-like CD8(+) T Cells Sustain the Immune Response to Chronic Viral Infections. Immunity 45, 415–427 (2016). 10.1016/j.immuni.2016.07.021

55 Im, S. J. et al. Defining CD8+ T cells that provide the proliferative burst after PD-1 therapy. Nature 537, 417–421 (2016). 10.1038/nature19330

56 Khan, O. et al. TOX transcriptionally and epigenetically programs CD8(+) T cell exhaustion. Nature 571, 211–218 (2019). 10.1038/s41586-019-1325-x

57 Miller, B. C. et al. Subsets of exhausted CD8(+) T cells differentially mediate tumor control and respond to checkpoint blockade. Nat Immunol 20, 326–336 (2019). 10.1038/s41590-019-0312-6

58 Street, K. et al. Slingshot: cell lineage and pseudotime inference for single-cell transcriptomics. BMC Genomics 19, 477 (2018). 10.1186/s12864-018-4772-0

59 Xie, F. et al. Breast cancer cell-derived extracellular vesicles promote CD8(+) T cell exhaustion via TGF-beta type II receptor signaling. Nat Commun 13, 4461 (2022). 10.1038/s41467-022-31250-2

60 Hu, Y. et al. TGF-beta regulates the stem-like state of PD-1+ TCF-1+ virus-specific CD8 T cells during chronic infection. J Exp Med 219 (2022). 10.1084/jem.20211574

61 Thomas, D. A. & Massague, J. TGF-beta directly targets cytotoxic T cell functions during tumor evasion of immune surveillance. Cancer Cell 8, 369–380 (2005). 10.1016/j.ccr.2005.10.012

62 Shlien, A. & Malkin, D. Copy number variations and cancer. Genome Med 1, 62 (2009). 10.1186/gm62

63 Gomez-Abad, C. et al. PIM2 inhibition as a rational therapeutic approach in B-cell lymphoma. Blood 118, 5517–5527 (2011). 10.1182/blood-2011-03-344374

64 Nann, D. et al. Follicular lymphoma t(14;18)-negative is genetically a heterogeneous disease. Blood Adv 4, 5652–5665 (2020). 10.1182/bloodadvances.2020002944

65 Eide, M. B. et al. Genomic alterations reveal potential for higher grade transformation in follicular lymphoma and confirm parallel evolution of tumor cell clones. Blood 116, 1489–1497 (2010). 10.1182/blood-2010-03-272278

66 Ma, M. C. J. et al. Subtype-specific and co-occurring genetic alterations in B-cell non-Hodgkin lymphoma. Haematologica 107, 690–701 (2022). 10.3324/haematol.2020.274258

67 Kim, M. et al. Clinicopathological and prognostic significance of BCL2, BCL6, MYC, and IRF4 copy number gains and translocations in follicular lymphoma: a study by FISH analysis. Leuk Lymphoma 61, 3342–3350 (2020). 10.1080/10428194.2020.1815017

68 O’Shea, D. et al. The presence of TP53 mutation at diagnosis of follicular lymphoma identifies a high-risk group of patients with shortened time to disease progression and poorer overall survival. Blood 112, 3126–3129 (2008). 10.1182/blood-2008-05-154013

69 Weigert, O. et al. Molecular ontogeny of donor-derived follicular lymphomas occurring after hematopoietic cell transplantation. Cancer Discov 2, 47–55 (2012). 10.1158/2159-8290.CD-11-0208

70 He, R. et al. Follicular CXCR5-expressing CD8(+) T cells curtail chronic viral infection. Nature 537, 412–428 (2016). 10.1038/nature19317

71 Leong, Y. A. et al. CXCR5(+) follicular cytotoxic T cells control viral infection in B cell follicles. Nat Immunol 17, 1187–1196 (2016). 10.1038/ni.3543

72 Chu, F. et al. CXCR5(+)CD8(+) T cells are a distinct functional subset with an antitumor activity. Leukemia 33, 2640–2653 (2019). 10.1038/s41375-019-0464-2

73 Wahlin, B. E., Sander, B., Christensson, B. & Kimby, E. CD8+ T-cell content in diagnostic lymph nodes measured by flow cytometry is a predictor of survival in follicular lymphoma. Clin Cancer Res 13, 388–397 (2007). 10.1158/1078-0432.CCR-06-1734

74 Collins, D. R. et al. Cytolytic CD8(+) T cells infiltrate germinal centers to limit ongoing HIV replication in spontaneous controller lymph nodes. Sci Immunol 8, eade5872 (2023). 10.1126/sciimmunol.ade5872

75 Dunn, G. P., Old, L. J. & Schreiber, R. D. The three Es of cancer immunoediting. Annu Rev Immunol 22, 329–360 (2004). 10.1146/annurev.immunol.22.012703.104803

76 Dunn, G. P., Old, L. J. & Schreiber, R. D. The immunobiology of cancer immunosurveillance and immunoediting. Immunity 21, 137–148 (2004). 10.1016/j.immuni.2004.07.017

77 Gravelle, P. et al. Impaired functional responses in follicular lymphoma CD8(+)TIM-3(+) T lymphocytes following TCR engagement. Oncoimmunology 5, e1224044 (2016). 10.1080/2162402X.2016.1224044

78 Han, G. et al. Follicular Lymphoma Microenvironment Characteristics Associated with Tumor Cell Mutations and MHC Class II Expression. Blood Cancer Discov 3, 428–443 (2022). 10.1158/2643-3230.BCD-21-0075

79 Li, J. et al. Loss of CREBBP and KMT2D cooperate to accelerate lymphomagenesis and shape the lymphoma immune microenvironment. Nat Commun 15, 2879 (2024). 10.1038/s41467-024-47012-1

80 Mariathasan, S. et al. TGFbeta attenuates tumour response to PD-L1 blockade by contributing to exclusion of T cells. Nature 554, 544–548 (2018). 10.1038/nature25501

81 Tauriello, D. V. F. et al. TGFbeta drives immune evasion in genetically reconstituted colon cancer metastasis. Nature 554, 538–543 (2018). 10.1038/nature25492

82 Peranzoni, E. et al. Macrophages impede CD8 T cells from reaching tumor cells and limit the efficacy of anti-PD-1 treatment. Proc Natl Acad Sci U S A 115, E4041–E4050 (2018). 10.1073/pnas.1720948115

83 Smith, A. et al. Lymphoma incidence, survival and prevalence 2004-2014: sub-type analyses from the UK’s Haematological Malignancy Research Network. Br J Cancer 112, 1575–1584 (2015). 10.1038/bjc.2015.94

84 Ren, P., Dong, X. & Vijg, J. Age-related somatic mutation burden in human tissues. Front Aging 3, 1018119 (2022). 10.3389/fragi.2022.1018119

85 Lancaster, J. N. Aging of lymphoid stromal architecture impacts immune responses. Semin Immunol 70, 101817 (2023). 10.1016/j.smim.2023.101817

86 Cinti, I. & Denton, A. E. Lymphoid stromal cells-more than just a highway to humoral immunity. Oxf Open Immunol 2, iqab011 (2021). 10.1093/oxfimm/iqab011

87 Spira, A. et al. Leveraging premalignant biology for immune-based cancer prevention. Proc Natl Acad Sci U S A 113, 10750–10758 (2016). 10.1073/pnas.1608077113

88 Spira, A. et al. Precancer Atlas to Drive Precision Prevention Trials. Cancer Res 77, 1510–1541 (2017). 10.1158/0008-5472.CAN-16-2346

89 Stanton, S. E., Castle, P. E., Finn, O. J., Sei, S. & Emens, L. A. Advances and challenges in cancer immunoprevention and immune interception. J Immunother Cancer 12 (2024). 10.1136/jitc-2023-007815

90 Brink, R. et al. The SW(HEL) system for high-resolution analysis of in vivo antigen-specific T-dependent B cell responses. Methods Mol Biol 1291, 103–123 (2015). 10.1007/978-1-4939-2498-1_9

91 Goodnow, C. C. et al. Altered immunoglobulin expression and functional silencing of self-reactive B lymphocytes in transgenic mice. Nature 334, 676–682 (1988). 10.1038/334676a0

